# Single-cell RNA-seq reveals dynamic transcriptome profiling in human early neural differentiation

**DOI:** 10.1101/384131

**Authors:** Zhouchun Shang, Dongsheng Chen, Quanlei Wang, Shengpeng Wang, Qiuting Deng, Liang Wu, Chuanyu Liu, Xiangning Ding, Shiyou Wang, Jixing Zhong, Doudou Zhang, Xiaodong Cai, Shida Zhu, Huanming Yang, Longqi Liu, J. Lynn Fink, Fang Chen, Xiaoqing Liu, Zhengliang Gao, Xun Xu

## Abstract

**Background:** Investigating cell fate decision and subpopulation specification in the context of the neural lineage is fundamental to understanding neurogenesis and neurodegenerative diseases. The differentiation process of neural-tube-like rosettes *in vitro* is representative of neural tube structures, which are composed of radially organized, columnar epithelial cells and give rise to functional neural cells. However, the underlying regulatory network of cell fate commitment during early neural differentiation remains elusive.

**Results:** In this study, we investigated the genome-wide transcriptome profile of single cells from six consecutive reprogramming and neural differentiation time points and identified cellular subpopulations present at each differentiation stage. Based on the inferred reconstructed trajectory and the characteristics of subpopulations contributing the most towards commitment to the central nervous system (CNS) lineage at each stage during differentiation, we identified putative novel transcription factors in regulating neural differentiation. In addition, we dissected the dynamics of chromatin accessibility at the neural differentiation stages and revealed active c/s-regulatory elements for transcription factors known to have a key role in neural differentiation as well as for those that we suggest are also involved. Further, communication network analysis demonstrated that cellular interactions most frequently occurred among embryoid body (EB) stage and each cell subpopulation possessed a distinctive spectrum of ligands and receptors associated with neural differentiation which could reflect the identity of each subpopulation.

**Conclusions:** Our study provides a comprehensive and integrative study of the transcriptomics and epigenetics of human early neural differentiation, which paves the way for a deeper understanding of the regulatory mechanisms driving the differentiation of the neural lineage.

## Background

The nervous system contains complex molecular circuitry in developmental processes. In humans, there is a paucity of data describing early neural development and the corresponding cellular heterogeneity at various stages. To our knowledge, neural tube formation and closure is crucial for embryonic central nervous system (CNS) development and the process of neurulation. Previous studies have reported that neural tube closure is strongly controlled by both genetic and epigenetic factors and is sensitive to environmental influences [1–3]. Perturbations in this delicately balanced and orchestrated process can result in neural tube defects (NTDs) giving rise to birth defects such as spina bifida, anencephaly and encephaloceles. However, the formation and closure of the neural tube *in vivo* during week 3 and 4 of human gestation is a transient event and is therefore difficult to capture. Moreover, the limited accessibility of human abortive fetuses at such an early stage precludes a thorough investigation of human early neural development.

Human pluripotent stem cells (hPSCs), including embryonic stem cells (ESCs) and induced pluripotent stem cells (iPSCs), can be differentiated into all cell types, including neural cells, offering a promising *in vitro* model for tracing early cell lineages and studying the cell fate specification of human neural differentiation [4, 5]. Previous studies have indicated that inhibition of bone morphogenetic protein (BMP) signalling or activation of fibroblast growth factor (FGF) signalling is needed for induction of the neuroectoderm from ESCs [6, 7]. A striking feature of differentiating stem cells *in vitro* is that they form neural tube-like rosettes which are composed of radially organized columnar epithelial cells that resemble the process of neurulation. The progenitor cells in rosettes gradually give rise to functional cells (e.g., more restricted progenitors and neuronal precursors, mimicking the process of neurulation and neural tube growth) which represent neural tube structures [8]. These cellular processes suggest that distinct cell fate decisions and lineage commitments occur during rosette formation. However, the corresponding underlying mechanisms of the regulation of cell fate commitment during early neural differentiation remain largely unknown.

The advance of single cell trans-omics technology has offered incisive tools for revealing heterogeneous cellular contexts and developmental processes [9–11]. Single cell RNA-seq (scRNA-seq) has been applied to the study of cellular heterogeneity as well as to the identification of novel subtypes or intermediate cell groups in multiple contexts [12–15], and may help delineate unexpected features of neural developmental biology and facilitate the study of cellular states and neurogenesis processes. In the present study, we used scRNA-seq and ATAC-seq (assay for transposase-accessible chromatin using sequencing) to investigate human early neural differentiation. Our analysis reveals the landscape of the transcriptome and c/s-regulatory elements during this process and creates an unbiased classification of cell subpopulations during differentiation, providing a comprehensive description of transcriptomic and epigenetic patterns in cell fate decision. The differentiation system of hiPSCs provides access to the very early stage of neural development and may serve as a source of specialized cells for regenerative medicine as well as supporting further investigations of neural tube defects.

## Data description

Here, we applied a well-adopted neural induction protocol and generated neural progenitor cells (NPCs) by forming neural rosettes */n v/tro* [8, 16]. We analysed several different differentiation stages of cells, including hiPSCs, embryoid body (EB), early rosettes (hereafter termed Ros-E, post-3 days of rosettes formation), late rosettes (hereafter termed Ros-L, post-5 days of rosettes formation), NPCs, and the original somatic fibroblasts (Fib). scRNA-seq was performed at discrete time points (e.g., Fib, iPSCs, EB, Ros-E, Ros-L and NPCs), and we captured 96, 80, 81, 82, 93, and 95 single cells, respectively, for each stage with the purpose of studying differentiation transition events. The quality of sequencing data was evaluated and filtered by a quality control (QC) pipeline developed in-house (see Methods for details). In addition, bulk ATAC-seq with two biological replicates was applied to the cell stages iPSCs, EB, Ros-E, Ros-L and NPCs to measure the regulome dynamics during neural differentiation (fig. 1a).

**Fig. 1.**
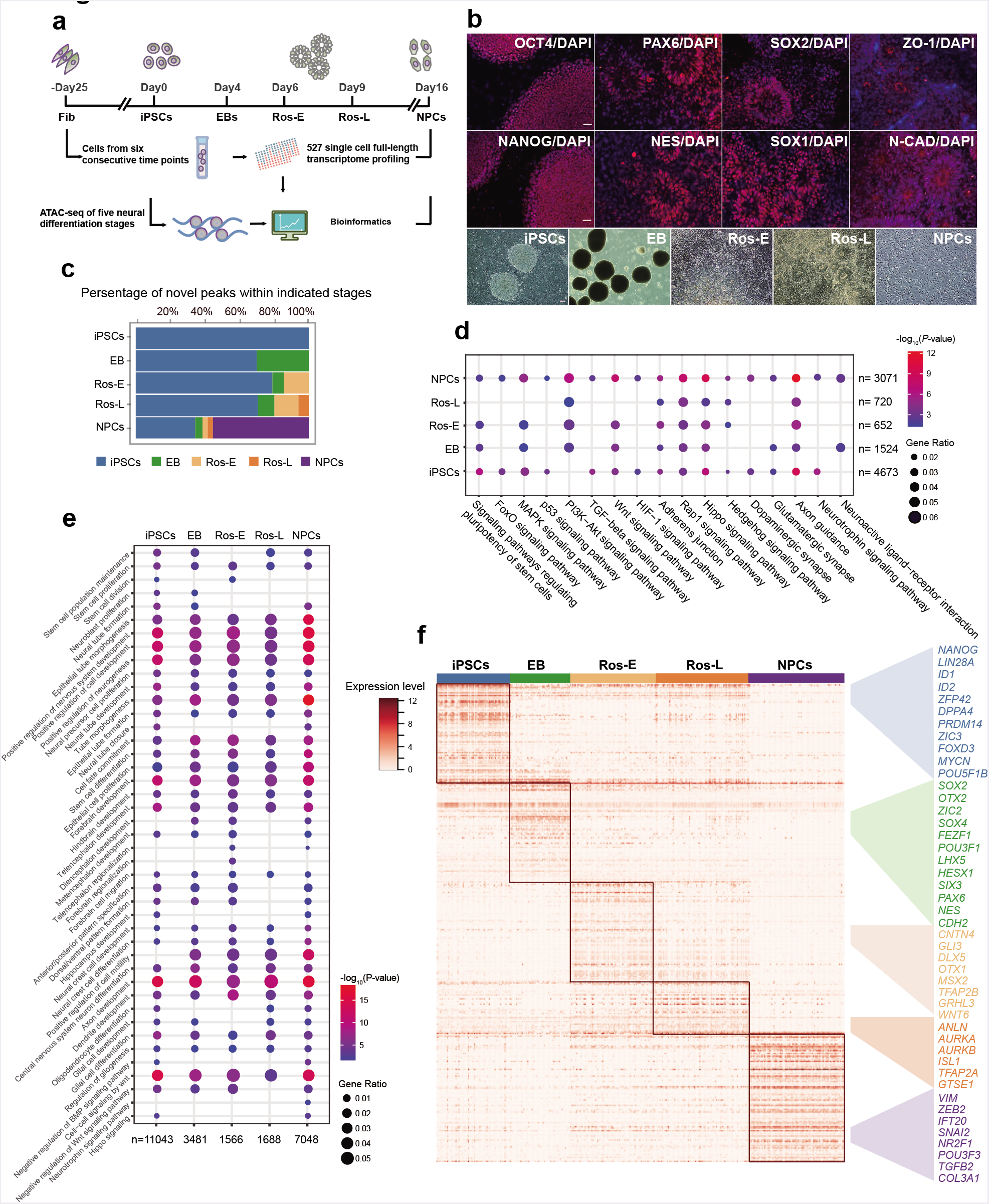
Transcriptome and regulome dynamics during human early neural differentiation. **a** Schematic illustration of experimental strategy. **b** Bright field and immunostaining of well-defined markers for iPSCs including OCT4 and NANOG, and for neural rosettes including PAX6, NES (NESTIN), SOX2, SOX1, ZO-1 and N-CAD (N-CADHERIN, also known as CDH2). Scale bar represents 50 pm. **c** Dynamic distribution of novel peaks (active c/s-regulatory elements) within indicated cell stages. **d** KEGG enrichment analysis of novel peaks within each cell stage as indicated respectively. **e** GO term annotation of novel peaks within each cell stage as indicated respectively. **f** Stage specific genes highlight with color specific to the respective neural differentiation cell stage (adjusted *P*-value < 0.01).

## Analyses

### Differential transcriptome and regulome dynamics throughout human early neural differentiation

Since the development of human ESCs and iPSCs, the ability to investigate human neurogenesis and neurological diseases via an *in vitro* differentiation model has vastly improved [4, 17]. Subsequently, artificial neural cells have been successfully generated using a variety of protocols by several laboratories [18–23]. Here, we followed a well-adopted neural induction protocol and generated NPCs by forming neural rosettes via inhibition of TGFβ, AMPK and BMP signalling pathways and activation of the FGF signalling pathway [8, 16]. We analysed different differentiation stages of the cells including iPSCs, EB, Ros-E, Ros-L, and NPCs as well as the original somatic fibroblasts (Fib). The iPSC aggregates were induced to neuroepithelial cells (NE) and followed by neural tube-like rosettes formation (fig. 1b). Firstly, pluripotency-associated transcription factors (TFs) (e.g., OCT4, NANOG) were significantly expressed in hiPSCs, suggesting that these cells did exhibit a stem cell phenotype. The subsequent formation of neural rosettes was confirmed by morphology, apical localization of ZO-1, a tight junction protein, and co-localisation of the neuroepithelial marker N-CADHERIN (N-CAD, also known as CDH2) at the junctions. Additional neural markers such as PAX6, NESTIN, SOX2, and SOX1 were also found to be highly enriched in the rosette stage (fig. 1b).

Cell stages are usually determined by a complement of TFs or master regulators which regulate hundreds of genes associated with various cellular functions. To study the genomic features associated with open chromatin regions, we classified ATAC peaks based on the location of the peak centre. More than 16,000 peaks were identified for each cell stage (Additional file 1: Figure S1a) with the majority located in introns and enhancers/promoters, genomic regions that are known to harbour a variety of c/s-regulatory elements and are subjected to regulation by TFs (Additional file 1: Figure S1b). Furthermore, we observed that ATAC peaks were significantly enriched at regions near transcription start sites (TSS) (Additional file 1: Figure S1c). These observations were reproducible across two replicates with a very high Pearson correlation coefficient (>=0.954) (Additional file 1: Figure S11, d).

It is widely reported that chromatin structures undergo widespread reprogramming during cell status transition, with some genomic regions becoming compacted or opened, leading to the switching on or off of a repertoire of genes responsible for cell fate decision [24–29]. We studied the dynamic chromatin landscape by tracing the temporal origins of ATAC peaks at each stage with peaks non-overlapping with existing ones that were annotated as novel peaks. We assumed that those peaks, conserved among differentiation stages, are associated with housekeeping genes while stage-dynamic peaks are likely to represent *c/s*-regulatory elements important for cell status transition. As expected, we observed the introduction of roughly 10-50% of novel peaks in each stage, accompanied by the disappearance of several pre-existing ATAC peaks. Notably, more novel peaks appeared at the NPCs stage than at other stage (fig. 1c). GO term analysis of genes residing in novel peaks across the differentiation stages showed enrichment of “axon development”, “positive regulation of nervous system development”, “epithelial tube morphogenesis”, “positive regulation of neurogenesis”, “cell-cell signalling by Wnt”, “forebrain development”, “hindbrain development”, “telencephalon development”, “neural precursor cell proliferation”, and “cell fate commitment”. “Neurotrophin signalling pathway” was also found to be enriched, but was specifically associated with NPCs. KEGG enrichment analysis showed that “FoxO signalling pathway”, a pathway which is known to play an important role in NPC proliferation, and “neuroactive ligand-receptor interaction” were enriched in NPCs stage (Fig. 1d, e), suggesting that specific c/s-regulatory elements regulating neural differentiation are being staged (poised) for stem cell fate specification and conversion.

To reveal the detail of chromatin accessibility dynamics during neural differentiation, we also analysed the gained or lost peaks at each stage compared with the previously neighbouring one. We observed that the number of gained peaks was with the largest increase at the NPCs stage while the number of lost peaks was relatively high at Ros-E stage (Additional file 2: Figure S2a). Next, we studied the genomic distribution of these dynamic peaks and found that both the gained and lost peaks were located mostly in distal intergenic regions and promoter regions (Additional file 2: Figure S2b). This observation indicates that distal and promoter regions are more dynamic compared to other genomic regions during neural differentiation process.

To gain insight into the potential function of closing (lost) peaks dynamics, we carried out GO enrichment analysis on the genes associated with lost peaks at each stage. The GO terms analysis showed that “mesoderm morphogenesis”, “endoderm development”, “gastrulation” and “nodal signalling pathway” were solely enriched at EB stage, indicating that upstream, as well as other lineage development, was relatively repressed by closing related cis-regulatory regions. Other cell fate conversion terms such as “neural crest cell differentiation”, “osteoclast differentiation”, and “regulation of cartilage development” were enriched at Ros-E stage, together with the annotation results of novel peaks, indicating that the chromatin accessibility prepared for the neural lineage conversion by opening/closing up specific *cis*-regulatory regions which facilitated the neural transition cascades (Fig. 1d, e and Additional file 2: Figure S2d, e).

Furthermore, we identified stage-specific peaks at iPSCs, EB, Ros-E, Ros-L and NPCs using motif enrichment analysis (see Methods). Further GO term and KEGG enrichment analysis showed very similar results with annotation analysis of novel peaks in corresponding cell stages (Additional file 3: Figure S3). These findings strongly suggest that the novel, gained and lost, as well as stage-specific peaks, represent cell status and cell fate transitions that progress neural differentiation and that the landscape of c/s-regulatory element accessibility throughout the differentiation process is highly dynamic.

To more thoroughly investigate the molecular mechanisms governing neural differentiation we profiled the transcriptomes of 527 single cells. Single cell RNA-seq libraries were generated using Smart-Seq2 method [30], followed by sequencing approximately 6 million reads per cell. For subsequent analysis, we focused on 445 cells that passed the quality control (QC, Methods, Additional file 4: Figure S4a, b) and ERCC correlation filter (Methods, Additional file 4: Figure S4c). 7003 to 8560 expressed genes were detected per cell (Additional file 4: Figure S4d), including TFs that were relatively highly expressed at the EB and NPCs stages, while, intriguingly, pseudogenes were relatively highly expressed at the Ros-E and NPCs stages (Additional file 4: Figure S4e). We also identified a variety of genes: 3524, 3855, 2023, 1804 and 6211 specifically expressed at the iPSCs, EB, Ros-E, Ros-L and NPCs stages, respectively (Additional file 4: Figure S4f). Many of these stage-specific genes include some well-known pluripotent genes *(NANOG, ID1, ID2, ZFP42, LIN28A, DPPA4);* early neural markers *(SOX2, OTX2, OTX1, PAX6);* and genes that both regulate neural development and are critical to proliferative NPCs *(SOX4, SIX3, CDH2, ZIC2)* (Fig. 1f and Additional file 4: Figure S4h).

Because the neural rosette recapitulates neural tube development *in vitro*, we paid particular attention to the Ros-E and Ros-L stages. Unsurprisingly, a large proportion of up-regulated genes in the Ros-E stage were associated with nervous system development including *TFAP2A, CNTN4, GLI3, DLX5* and *OTX1)* (fig. 1f). Of particular interest is the gene *GRHL3.* Expression of this gene is associated with neural tube closure in mice [31, 32] and we observed this gene to be highly expressed at Ros-E in human cells, suggesting that its role in neural tube closure may be conserved across mammals or possibly chordates. *TFAP2A* (transcription factor AP-2 alpha) and *TFAP2B* (transcription factor AP-2 beta) have been proposed as master regulators of the neural crest cell and loss of function of transcription factor AP-2 in mice is strongly associated with a cranial neural tube defect phenotype [33]. In our system, *TFAP2B* and *TFAP2A* were relatively highly expressed at both the Ros-E and -L stages, suggesting transcription factor AP-2 may coordinate the specialized distal *cis*-regulatory elements for downstream regulations in human. We also observed expression of *ANLN* (Anillin actin binding protein) at the Ros-L stage, suggesting that neuronal migration and neurite growth might occur by the linking of RhoG to the actin cytoskeleton in neural rosettes [34]. Similarly, our data showed that *AURKA* (aurora kinase A) and *AURKB* (aurora kinase B) were both expressed at the Ros-L stage, echoing previous findings that the aPKC–Aurora A–NDEL1 pathway plays an essential role in neurite elongation through modulating microtubule dynamics [35]. Finally, the neuron fate commitment protein, *TGFB2*, the nervous system development regulator, *ZEB2*, and the neural precursor cell proliferation-associated protein, *IFT20*, were enriched at NPCs stage (fig. 1f).

An unexpected finding was that some of the most important neural TFs exhibited heterogeneous expression within the same cell stage (e.g., *ZIC2, OTX2, HESX1, DLX3, LHX5)* (Fig. 1f and Additional file 4: Figure S4h). This inspired us to dissect the subpopulations of cells within each cell stage to better understand the significance of this result.

### Heterogeneous cellular subpopulations were identified at each developmental stage

To evaluate the overall distribution of cells at each of the six stages during reprogramming and neural differentiation, we first performed an unsupervised analysis using all expressed genes (QC, see Methods) as input to t-distributed stochastic neighbour embedding (t-SNE) for visualization. This analysis showed distinct clusters for each differentiation stage, supporting our observation of heterogeneous gene expression during these stages (fig. 2a). Because previous studies have showed that TFs and c/s-regulatory elements are highly informative in reflecting cell identity [36], we used a machine classifier to determine the subsets of TFs that best clustered cells into putative cell populations. We were then able to identify distinct subpopulations at each cell stage (Fib1, Fib2, EB1, EB2, EB3, Ros-E1, Ros-E2, Ros-L1, Ros-L2, Ros-L3, NPC1, NPC2 and NPC3) (Methods, Fig. 2, Additional file 5-8: Figure S5-8). As we found no remarkable differential expression of pluripotency-associated genes (e.g., *NANOG, ID1, ID2, LIN28A, SOX2, DPPA4, ZFP42, TRIM28)* at the iPSCs stage (Additional file 4: Figure S4g), we did not include iPSCs in the following analyses.

**Fig. 2.**
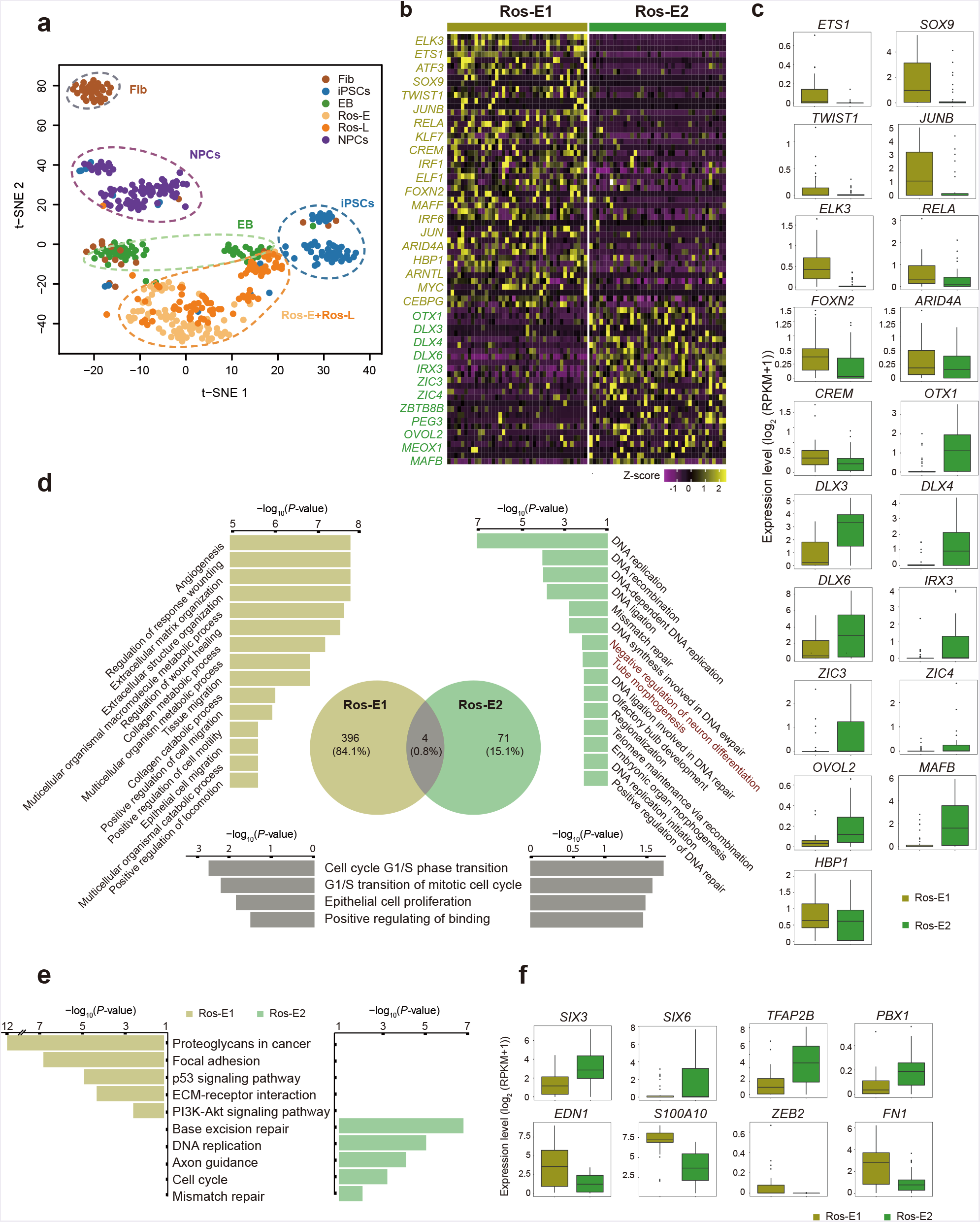
Cell heterogeneity and identification of subsets within Ros-E stage. **a**T-SNE analysis of different cell stages as indicated with different color (n = 445). Number of successfully profiled single cells per cell stage: Fib (n = 54); iPSCs (n = 71); EB (n = 57); Ros-E (n = 81); Ros-L (n = 92); NPCs (n = 90). Each dot represents an individual cell. b Heatmap shows scaled expression [log_2_ (RPKM+1)] of discriminative TF sets for each cluster at Ros-E stage, *P*-value ≤ 0.01. Color scheme is based on z-score distribution from −1 (purple) to 2 (yellow). c Box plot of discriminative TFs for specific subpopulation at Ros-E stage. d GO term enrichment of differentially up-regulated genes respective to indicated subpopulation (highlighted with color: Ros-E1 is yellow; Ros-E2 is green; overlapped GO terms of Ros-E1 and Ros-E2 are grey). e Top 5 differential pathway in Ros-E1 and Ros-E2 respectively by KEGG enrichment analysis. f Representative box plots of subpopulation specific genes identified by SCDE (single-cell differential expression), adjusted *P*-value < 0.01.

### Fibroblasts (Fib) stage

Fibroblasts (Fib) are a very well-adopted original somatic cell resource for iPSCs reprogramming; many direct conversions from fibroblast to functional neurons have been reported [37, 38]. Here, we dissected two subpopulations of human dermal fibroblasts (Fib1 and Fib2) with distinct molecular features, showing significantly higher expression of several important pluripotency- and neural-associated transcription factors such as *SOX2, LIN28, SOX11, ZIC2, FEZF1* and *SIX3* in Fib2 (Additional file 5: Figure S55, a). GO terms identified by up-regulated genes between the two subsets showed “chromosome segregation”, “positive regulation of nervous system development”, “stem cell population maintenance”, “positive regulation of cell cycle”, “neural precursor cell proliferation” and “chromatin remodeling” as solely enriched in the Fib2 subpopulation (Additional file 5: Figure S5c). KEGG enrichment analysis showed “cell cycle” term was specifically associated with the Fib2 subset (Additional file 5: Figure S5d). Furthermore, we observed that fibroblasts were distributed into two distinct groups called Fib-Group1 and Fib-Group2 based on their location in the Fig. 2a. Of note, the majority of cells in Fib-Group1 and Fib-Groups2 were composed of Fib1 and Fib2, respectively. Moreover, cells from Fib2 subset clustered together with EB cells (Additional file 5: Figure S5e). Together with the molecular features of Fib2 subset (Additional file 5: Figure S5b), we proposed Fib2 subset might possess high potential for iPSCs reprogramming and neural conversion. Thus, based on the differentially expressed genes and CD markers dataset (HUGO Gene Nomenclature Committee, HGNC), we further inferred several cell surface markers of Fib2 (e.g., *FGFR2, F11R, PROM1, BST2, ITGA6* and *EPCAM)* although these surface markers showed heterogeneously expressed levels within the Fib2 subset (Additional file 5: Figure S5f).

### Embryoid body (EB) stage

For the three EB subpopulations (EB1, EB2 and EB3), we identified genes that were up-regulated compared to the iPSCs stage, respectively. These genes were enriched in “fetal brain cortex”, “epithelium” and “brain” terms by DAVID using tissue enrichment analysis (Additional file 6: Figure S6a) which suggests that the biological processes of brain development and neural differentiation initiation are occurring during the iPSCs-to-EB stage transition and these processes are shared by each EB subpopulation. Moreover, most neural TFs and cell-specific markers were expressed commonly among EB subpopulations (e.g., *SOX2, ZIC2, SOX11, SOX4, SIX3)* (Additional file 6: Figure S6b) and some of these TFs play a crucial role in neural tube formation. However, some important neural TFs, such as *FOXO1* and *FOXO3*, which play an important role in NPC proliferation and self-renewal [39]; *TULP3*, which regulates the SHH signalling pathway and modulates neural tube development [40]; and *POU2F1*, which regulates *NESTIN* gene expression during P19 cell neural differentiation and CNS development [41], showed significantly high expression in the EB3 subpopulation, but low expression in the EB1 and EB2 subpopulations (Additional file 6: Figure S66, c). This suggests that different subpopulations contain specific molecular signatures and different differentiation states or potentials.

### Early rosette (Ros-E) stage

During the Ros-E stage, which is composed of NE and the cells in the early stage of rosette formation, we observed expression of several master regulator genes associated with neural tube formation and closure including *SOX11, ZIC2, PAX3*, and *SNAI2* in both Ros-E subgroups (Ros-E1 and Ros-E2). However, genes involved in neural crest specifiers, such as *TWIST1* [42] and *SOX9*, which contribute to the induction and maintenance of neural stem cells and are enriched in neural crest cells [43–45]; and *ETS1*, which regulates neural crest development through mediating BMP signalling [46], were preferentially expressed in the Ros-E1 subpopulation (Fig. 2b, c). The ectoderm marker, *OTX1*, and genes involved in the ventral hindbrain marker (e.g., *IRX3)* were highly expressed in the Ros-E2 subgroup (Fig. 2b, c). GO term annotation analysis showed Ros-E1 and Ros-E2 shared GO terms of “cell cycle G1/S phase transition”, “G1/S transition of mitotic cell cycle”, “epithelial cell proliferation” and “positive regulating of binding” (fig. 2d) while “negative regulation of neuron differentiation” and “tube morphogenesis” were solely enriched in the Ros-E2 subpopulation (fig. 2d). KEGG enrichment analysis showed that “base excision repair”, “DNA replication”, “axon guidance”, “cell cycle” and “mismatch repair” were specifically associated with the Ros-E2 subset (fig. 2e). We further performed single-cell differential expression (SCDE) on both Ros-E subpopulations and identified additional differentially expressed genes between the two groups. *SIX3*, *SIX6*, *TFAP2B* and *PBX1* were more highly expressed in Ros-E2, whereas *EDN1, S100A10* and other genes related to neural crest migration, were highly expressed in Ros-E1 (fig. 2f).

### Late rosette (Ros-L) stage

At the Ros-L stage the genes *SNAI2, OTX2, FEZF1, ZIC3*, and *HESX1* showed significantly different expression patterns among the three distinguishable subpopulations (Ros-L1, Ros-L2 and Ros-L3) at the Ros-L stage (Additional file 7: Figure S7a, a). Moreover, *SMAD1* and *MYC*, two components in the Wnt signaling pathway which is critical for neural development [47, 48], were specifically enriched in the Ros-L3 subpopulation. Additionally, *JUNB* from the TGFβ signaling pathway was preferentially expressed in Ros-L3 compared to the other two subpopulations. Interestingly, *HAND1* and *ISL1*, which are mesoderm markers, and *TBX3*, which elicits endodermal determination, were highly expressed in the Ros-L1 subpopulation (Additional file 7: Figure S7a, a).

Of 648 GO terms identified by differentially expressed genes among these three subsets, 52 terms were shared by Ros-L1 and Ros-L3, such as “positive regulation of cell motility”, “angiogenesis”, “positive regulation of cellular component movement” and “epithelium migration” (Additional file 7: Figure S7c). A high proportion of cardiac development terms was enriched in Ros-L1, whereas DNA replication- and chromatin remodeling-related terms and pathways were significantly associated with Ros-L2. In addition, cell-substrate adhesion-related terms and cell cycle-related pathways were enriched in Ros-L3 (Additional file 7: Figure S7c, d).

Several subpopulation-specific genes were identified, including *NR2F1, ARID3A, SIX3, OTX2* and *FOXG1* at the NPCs stage (Additional file 8: Figure S8a, b). These observations suggest that significant TF expression patterns describe discrepant cell differentiation states or differentiation commitments inside the neural conversion process. Taken together, our results suggest that the subpopulation analyses accurately describe specific gene expression dynamics at each cell stage, which are likely masked in bulk sequencing analyses. Additionally, extrapolating from these observations, we can reason that reconstructing a differentiation trajectory based on the gene expression dynamics of individual subpopulations would allow us to dissect neural differentiation processes that we would otherwise be unable to observe.

### Tracking a reconstructed trajectory identifies key subpopulations during neural differentiation

Based on the subpopulations identified before, we wanted to track the gene expression dynamics of individual subpopulations to parse the neural differentiation processes and dissect the subpopulation with the highest contribution towards commitment to the CNS lineage. First, we reconstructed the differentiation trajectory using 8220 genes with variable expression. This showed that cells in stages from iPSCs to NPCs followed a sequential differentiation process where each stage exhibited a relatively discriminative region with some of the subpopulations overlapping (fig. 3a). Subsequently, based on the pairwise comparisons of TF expression levels, we inferred the connection of the subpopulations from the iPSCs stage to NPCs stage across the five-stage differentiation process (fig. 3b). TF expression levels were considered as strong indicators of cell stage and identity [36]. Here, we used the Pearson correlation coefficient to identify more biologically and molecularly similar cell subpopulations and considered them as cells within the same developmental linage [49]. As a result, iPSCs, EB3, Ros-E2, Ros-L3 and NPC1 were identified as the subpopulations contributing the most to commitment to the CNS lineage (fig. 3b). These findings were consistent with the specific gene expression pattern in individual subpopulations. For instance, *SOX13*, expressed in the developing nervous system and neural tube [50, 51], *FOXO1* [39] and *TULP3* [40] were significantly highly expressed in EB3 (Additional file 6: Figure S6c, d). *MAFB*, an important TF in hindbrain identity [52], was enriched in Ros-E2 (Fig. 2b, c); and other crucial neural development TFs, especially those involved in CNS development, such as *OTX1, DLX3, DLX6, ZIC3, ZIC4*, and *IRX3*, also showed high expression in the Ros-E2 subpopulation (Fig. 2b, c). Previously, we assumed that *GRHL3* might be involved in neural tube closure; here, the results showed that *GRHL3* was indeed significantly highly expressed in Ros-L3 (Additional file 7, Figure S7b). Additionally, neural crest regulators (e.g., *ETS1, ELK3, SOX9)* were enriched in Ros-L3 (Additional file 7, Figure S7b), suggesting that cell fate specification and differential cell status might exist even within subset. Strikingly, Ros-E2 and Ros-L3 that were identified in the dominant path to CNS lineage by correlation analysis were shown as a process of sequential conversion in our reconstructed trajectory (Fig. 3a, c). The molecular signature described by these subpopulations was consistent with the analysis that identified the key contributing subpopulations and encouraged us to perform additional cell fate decision analyses.

**Fig. 3.**
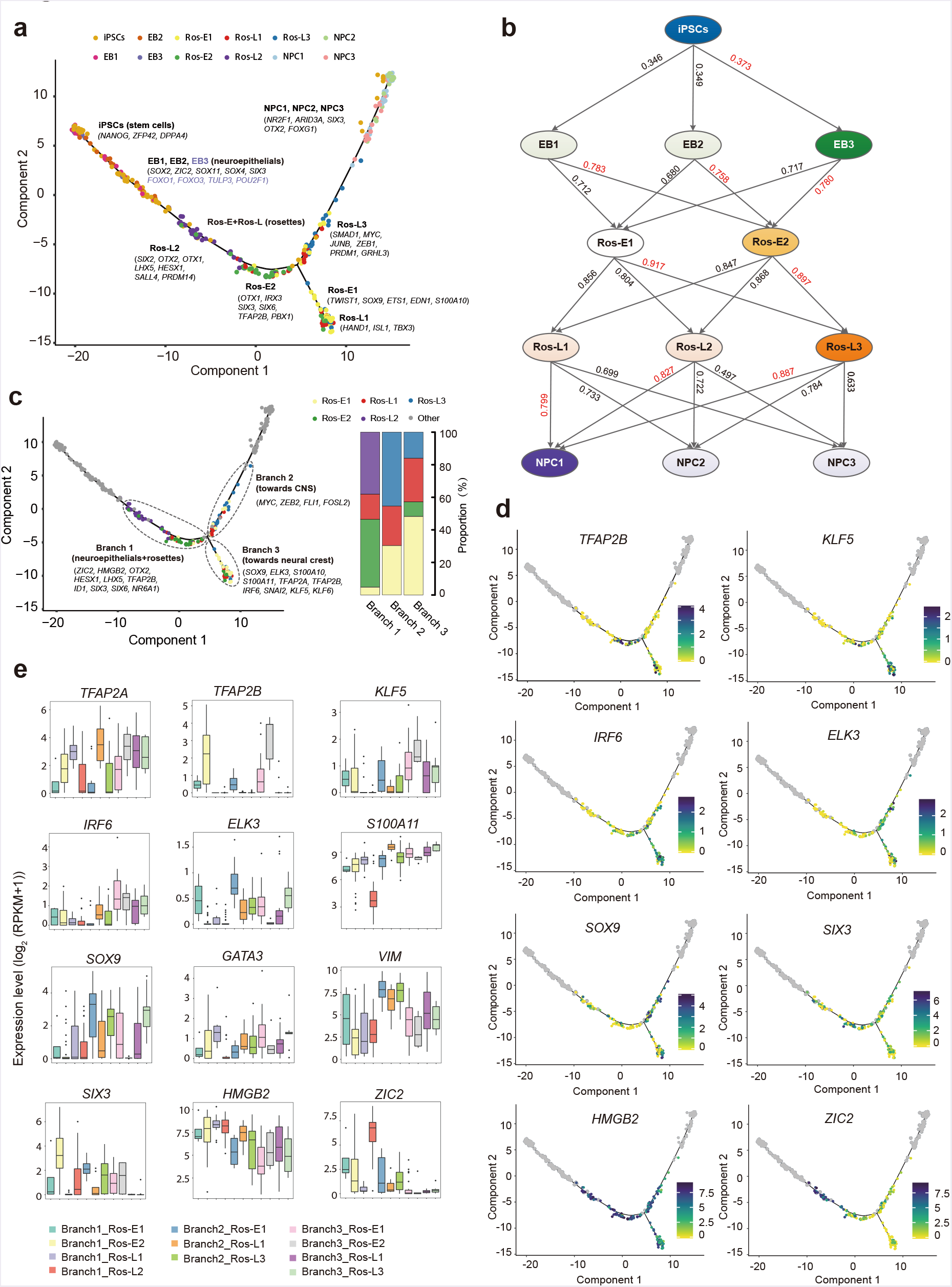
Cell fate specification revealed by reconstructed trajectory. **a**Differentiation trajectory constructed by 8220 variable genes across different cell stages. Selected marker genes specific to the respective cell stage/ subpopulation are indicated with black/purple color. b The connection of subpopulations from iPSCs to NPCs stage across the five-differentiation process identified by Pearson correlation coefficient. The Pearson correlation coefficient of the two comparisons is indicated on the arrow line, respectively. c The divarication point within rosette stage (Ros-E and Ros-L) across the differentiation trajectory, Branch 1, Branch 2 and Branch 3 based on their location on the differentiation trajectory are marked by dashed ellipse. Selected discriminative TFs specific to the respective branch are indicated. The columns represent the components of Branch 1, Branch 2 and Branch 3, respectively. d Expression pattern of selected differentially expressed TFs among the three branches on the reconstructed trajectory (adjusted P-value < 0.01). Color scheme is based on expression [log_2_ (RPKM+1)]. e Expression pattern of representative differentially expressed TFs across different components of the three branches.

Of note, there was a clear divarication within the rosette stages (Ros-E and Ros-L) across the differentiation trajectory, indicating cell fate decision might be made at this bifurcation point (fig. 3c). Here, we focused on the single cells in the rosette stages and called them Branch 1, Branch 2 and Branch 3 based on their location in the developmental trajectory (fig. 3c). Branch 3 was composed of Ros-E1 (n=27), Ros-L1 (n=15) and small proportion of Ros-E2 (n=5) and Ros-L3 (n=9, Fig. 3c). Previously, our observations showed that Ros-E1 was associated with neural crest cells (high expression of *TWIST1, SOX9, ETS1, EDN1* and *S100A10)* and Ros-L1 was likely related to mesoderm and endodermal determination (high expression of *HAND1, ISL1* and *TBX3)*, and these two subpopulations comprise the majority of cells in Branch 3. Further, we performed a pairwise comparison of gene expression across the three branches. The results showed that many neural TFs, such as markers of neural tube formation *(SOX4* and *SOX11);* the NSCs self-renewal and proliferation regulator *FOXO3;* and the NSC markers *NES, CDH2* and *FABP7*, were commonly expressed across all three branches, indicating the capacity for neural tube development and NSCs proliferation are a fundamental feature of neural rosettes (Additional file 9: Figure S9a, b). Strikingly, *ZIC2*, a member of the ZIC family of C2H2-type zinc finger proteins, associated with neural tube development [32], showed significantly low expression in Branch 3 (Fig. 3d, e). Some other neural development markers (e.g., *ZIC3, HMGB2, ID1, SIX3, SIX6, NR6A1)* were significantly lowly expressed in Branch 3 but highly expressed in Branch 1 (Fig. 3d, e, Additional file 9: Figure S9a, c). However, *TFAP2B*, encoding a member of the AP-2 family of TFs, and *ELK3*, essential for the progenitor progression to neural crest cell [53], was significantly highly expressed in Branch 3 but lowly expressed in Branch 2. Moreover, *SOX9, SNAI2, S100A11* and *TFAP2A*, previously shown to be highly expressed in neural crest cells [43,44,45,54], were markedly highly expressed in Branch 3, but not Branch 1 (Fig. 3d, e, Additional file 9: Figure S9a, c). *KLF5* and *IRF6* were significantly highly expressed in Branch 3 as well (Fig. 3d, e). These two TFs have been reported to be involved in phenotypic switching of vascular smooth muscle cells [55] and development of the palate in vertebrates involving cranial neural crest migration [56], respectively. These results indicate that cell fate specification might occur at the bifurcation point and, based on the observations, we speculate that Branch 1-to-Branch 2 has progressed more towards CNS and Branch 3 is probably composed of neural crest cells and other cells comprising this microenvironment.

### Construction of the TF regulatory network during cell status transition

To infer TFs which drive the progression of cell status from one stage to the neighbouring one, we performed SCDE analysis for those cell subpopulations committing to CNS lineage, resulting in 58, 123, 98 and 131 TFs differentially expressed among iPSCs vs EB3, EB3 vs Ros-E2, Ros-E2 vs Ros-L3, and Ros-L3 vs NPC1 comparisons (Additional file 10, 11: Figure S10, 11). Interestingly, *PRDM1*, which has been proposed to promote the cell fate specification RB sensory neurons in zebrafish [57], was significantly up-regulated from Ros-E2 to Ros-L3 (Additional file 10: Figure S10). In contrast, several well-characterized TFs were found to be significantly highly expressed in Ros-E2 (mainly resident in Branch 1) and down-regulated during the transition from early to late rosette development: *FOXG1*, cooperating with *Bmi-1* to maintain neural stem cell self-renewal in the forebrain; *MAFB*, the posterior CNS fate identifier and essential for hindbrain choroid plexus development [52, 58]; *DLX3* and *DLX5*, neural plate border specifier genes [58]; and *ID1*, a controller of stem cell proliferation during regenerative neurogenesis in the adult zebrafish telencephalon [59]. These results suggest that the expression patterns of neural-associated TFs undergo dramatic changes during neural differentiation with some TFs activated *(PRDM1*, etc.) and others repressed *(MAFB, FOXG1, ID1*, etc.) (Additional file 10: Figure S10). Furthermore, it was previously unknown that several of these TFs were involved in neural differentiation so our results have expanded the known biological functions of these molecules.

Among the 131 TFs exhibiting differential expression from Ros-L3 to NPC1, 80 TFs were up-regulated while 51 TFs were down-regulated (Additional file 11: Figure S11; Additional file 19: Table S1). Up-regulated TFs included *SNAI2*, a neural crest specifier [58]; *HIF1A*, required for neural stem cell maintenance and vascular stability in the adult mouse [60]; *SIX1*, which drives the neuronal developmental program in the mammalian inner ear [61]; *ETV1*, which orchestrates gene regulation during the terminal maturation program of cerebellar granule cells [62]; and *POU3F3*, which influences neurogenesis of upper-layer cells in the cerebral cortex [63] (Additional file 11: Figure S11). This is consistent with our previous observation that the main trajectory has progressed more towards to CNS. Of particular interest is *PRDM1*, whose expression increased from Ros-E2 to Ros-L3 and decreased during the progression from Ros-L3 to NPC1 (Additional file 10, 11: Figure S10, 11), suggesting that it might play multiple specific roles in neural differentiation.

Next, we inferred a regulatory network among those differentially expressed TFs based on known interactions collected in the STRING database [64]. Our results suggested that *SOX2* and *GATA3* were key regulators from iPSCs to EB3 (Additional file 12: Figure S12a); *TP53, SOX2, RELA, SIX3, ARNTL, ISL1, RARA, TP63*, GATA3, *SNAI2*, and *PAX3* were the key regulators from EB3 to Ros-E2 (Additional file 12: Figure S12b); *MYC, SOX2, PAX6, EGR1, PBX1, GLI3, PAX3, SIX3, FOXG1, OTX2, PAX7, PPARG, SOX9, MAFB, SIX6* and *ZIC1* were identified as key regulators from Ros-E2 to Ros-L3 (fig. 4a); and *SOX2, AR, MYCN, LEF1, PAX3, SNAI2, MSX1, SOX9, NR3C1, PARP1, RUNX1, EBF1, HIF1A, IRF6, IRF1, KLF5*, and *LIN28A* were predicted to be key regulators from Ros-L3 to NPC1 (fig. 4b).

**Fig. 4.**
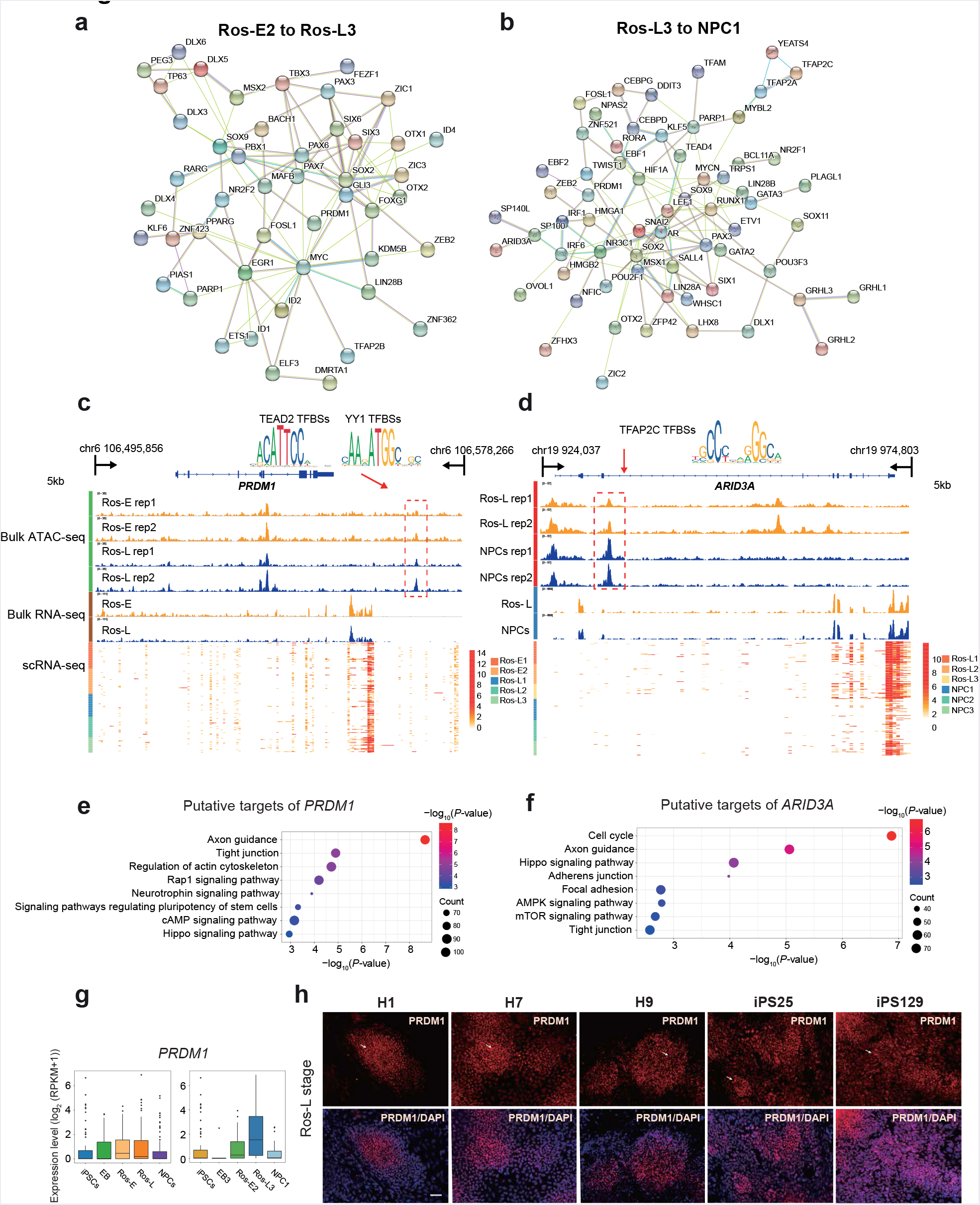
Key regulators and corresponding c/s-regulatory elements during neural differentiation. **a**Regulatory network of TFs differentially expressed between Ros-E2 and Ros-L3. b Regulatory network of differentially expressed TFs between Ros-L3 and NPC1. c, d IGV screenshots of ATAC-seq and bulk RNA-seq as well as the corresponding scRNA-seq heatmaps for putative neural regulator *PRDM1* (c) and *ARID3A* (d). Differential peaks in the dashed boxes possess putative TF motifs outlined in the form of sequence logo. e, f KEGG enrichment analysis of putative target genes under the regulation of *PRDM1* (e) and *ARID3A* (f). g Expression pattern of *PRDM1* at indicated cell stages (left) and subsets (right) during neural differentiation. h Immunostaining of PRDM1 at Ros-L stage across different genetic background cell lines (H1_ESCs, H7_ESCs, H9_ESCs, iPS25 and iPS129). Scale bar represents 50 pm.

To dissect the c/s-regulatory elements directing the expression of those regulators, we selected the differentially expressed TFs that showed differential ATAC peaks between neighbouring stages and performed motif scanning on the differential peaks. Focusing on the transition from Ros-E2 to Ros-L3, we found transcription factor binding sites (TFBSs) for TEAD2 and YY1 in a differential ATAC peak downstream of the *PRDM1* gene (fig. 4c). Multiple motifs for the transcription factor *TFAP2C* were found in a differential peak located in the intron of the *ARID3A* gene, which is a regulator responsible for the transition for Ros-L3 to NPCs (fig. 4d). Based on the temporal specificity of ATAC peaks and the existence of TF motifs in these regions, we propose that those elements are stage-specific *c/s*-regulatory elements regulating the expression of neural regulators in response to their upstream regulatory TFs.

To infer the putative targets of key regulators, we combined the information from ATAC peaks and motifs for TFs. All peaks containing motifs for a certain TF were annotated as TF-related peaks and genes proximal to the peak were considered as potential targets of that TF. Using these criteria, we predicted thousands of targets for each TF (Additional file 20: Table S2). To dissect the regulatory network of each TF, we conducted GO term and KEGG enrichment analysis for the putative target list of each key regulator. Our results suggested that, from Ros-E2 to Ros-L3, the targets for *PRDM1* were significantly enriched in pathways and GO terms associated with “axon guidance”, “hippo signalling pathway” and “neurotrophin signalling pathway” (Fig. 4e and Additional file 13: Figure S13). From Ros-L3 to NPC1, targets for *HIF1A, NR2F1, SOX9* and *TFAP2C* were enriched in KEGG pathways associated with “axon guidance” and “hippo signalling pathway” (Additional file 13: Figure S13). We further validated PRDM1 expression among different genetic background cell lines (H1_ESCs, H7_ESCs, H9_ESCs, iPS25 and iPSC129). The immunostaining showed that PRDM1 was expressed at Ros-L stage with heterogeneous expression level, though, the scRNA-seq data was not at a high level. Moreover, the results were uniformed across these cell lines (Fig. 4g, h).

### Inferring a cellular communication network among cell subpopulations within specific differentiation stages

Cell subpopulations with different functions are proposed to exhibit distinct expression profiles of ligands and receptors which primes cells for cell-type-specific interactions [65]. In this study, the cellular interactions were inferred using public ligand-receptor databases (see Methods). Briefly, 360, 182, 261 and 307 ligands/receptors were expressed within EB, Ros-E, Ros-L and NPCs subpopulations respectively, among which 304, 55, 124 and 162 interactions were identified within subpopulations at each differentiation time point (Fig. 5, Additional file 14-16: Figure S14-16 and Additional file 21: Table S3). The most frequent interactions were observed in the EB stage, implying that cells communicate extensively to coordinate differentiation programs during embryogenesis (Additional file 14: Figure S14). In contrast, much fewer interactions were predicted after the EB stage, suggesting communications decreased dramatically during the progression of lineage commitment. Notably, although comparable number of ligands and receptors were detected at EB (181 receptors and 179 ligands) and NPCs (128 receptors and 179 ligands) stage, only half the interactions (162) were inferred at NPCs stage compared to 304 ligand-receptor interactions at EB stage. (Additional file 14, 16: Figure S14, 16). The interactomes among Ros-L cells, with 31, 32 and 34 receptors from Ros-L1, Ros-L2 and Ros-L3 interacting with ligands from other cell subpopulations were inferred (fig. 5a). As expected, several interactions involving receptors and ligands previously known to play essential roles during neural development were identified in our study. For example, *WNT5A* and *EPHB6* were enriched in Ros-L1. *FZD5* and *LPAR4* were specifically expressed in Ros-L2. *PGF* and *ANGPT2* were up-regulated in Ros-L3 compared to other cell subpopulations (Fig. 5c, d, e). Overall, our study suggests that the specific expression spectrum of ligands and receptors and corresponding interactions can generally reflect the identity of cellular subpopulations.

**Fig. 5.**
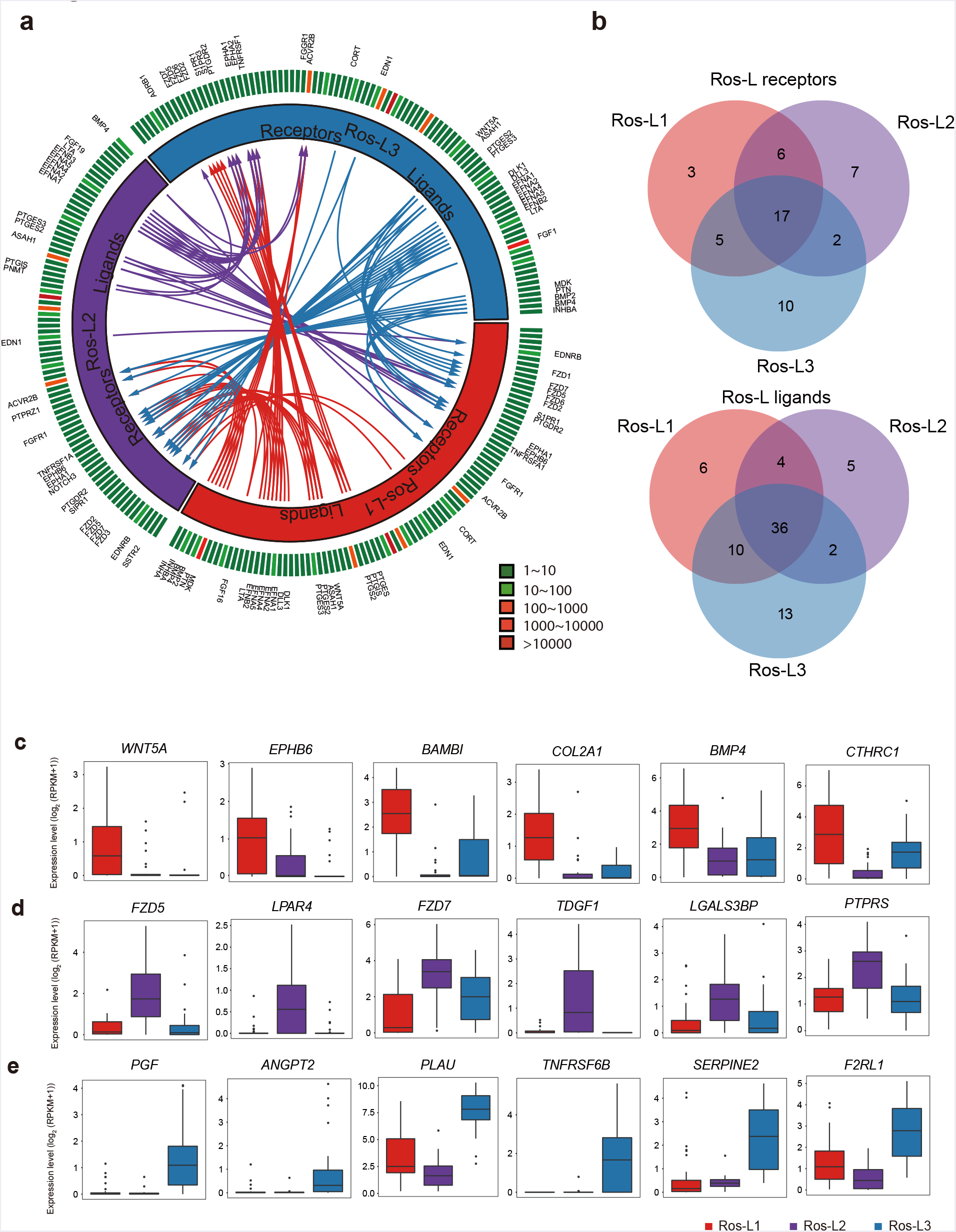
Putative receptor-ligand interactions in Ros-L subsets. **a**Putative signaling between expressed receptors and their ligands in Ros-L subsets. The inner layer compartments represent different cell subpopulations (Ros-L1, Ros-L2 and Ros-L3 were shown in red, purple and blue color respectively). The outer layer indicates the expression profiles of ligands and receptors expressed in each cell subset, with low expressed molecules in green color while high expressed ones in red color. Arrows indicate putative interactions between ligands and receptors among cell subsets. **b** Venn plot showing the overlapping of ligands and receptors among cellular subpopulations. **c, d, e** Expression level of receptors/ligands enriched in Ros-L1 (**c**), Ros-L2 (**d**) and Ros-L3 (**e**), respectively.

## Discussion

The regulation and molecular programs during embryonic neural development has long been investigated. However, much of this work has been limited to model organisms such as the mouse, zebrafish and *Drosophila* [36,40,56], due to the scarcity of human fetal tissue for research purposes. Our understanding of human early neural development, and particularly neural tube formation and the cell fate commitments of neural precursors in early stages, is still incomplete. To circumvent the challenges inherent in these investigations, namely the ability to study these processes *in vivo* in humans, we used hiPSCs and induced differentiation *in vitro* towards a neural cell fate using a well-established model. We characterised both the transcriptional profiles in single cells as well as chromatin accessibility at several critical stages during differentiation to inform this process at unprecedented resolution. This study has unveiled the dynamic transcriptome and regulome underlying the human early neural differentiation and identified functionally-distinct subpopulations within the various stages to have a more precise description of the factors defining the differentiation trajectory. Our analyses hint at the existence of a widespread regulatory network between TFs and their target genes, especially those associated with cellular reprogramming and differentiation. We were also able to construct minimal gene expression profiles based on ligands and receptors in each cell subpopulation which can be used to confidently infer cell identity.

During development *in vivo* the neuroectoderm folds to form the neural tube which is then patterned into regionally specialized subunits composed of progenitor cells. These cells subsequently give rise to regional progenies of neural cells [66]. There is some controversy in this field that formation of the EB would introduce *in vitro* culture variability in regional cells across different batches resulting in a relatively poor model of neural differentiation. The “dual-SMAD inhibition” method (inhibiting the SMAD-dependent TGFp and BMP signaling pathways) yielding neural epithelia in “monolayer culture” conditions [18] could alleviate the above concern. However, generation of neural rosette morphology */n v/tro* is considered equivalent to neural tube formation, recapitulating neural tube structure, which we believe is a promising research model for early neural differentiation. Neural differentiation of hiPSCs into NPCs starts with initial neural induction by appropriate dosages and gradients of many TFs and morphogenetic factors that are highly expressed in the developing brain. In this study, the induction cocktail used in the neural differentiation included SB431542, dorsomorphin, N2, B27, VEGF and bFGF supplemented at specific time points. The self-renewal program in human iPSCs is switched off and differentiation toward NE and NPCs is triggered [8, 16]. Previous results have shown that SB431542 enhances neural induction in EB derived from hESCs [65] by inhibiting the Lefty/Activin/TGFβ pathways and suppresses the mesodermal lineage (Brachyury) induction [18, 42]. Consistent with these previous studies, in our *m v/tro* system, treatment with SB431542, in combination with dorsomorphin, results in a dramatic decrease in *NANOG* expression and a concomitant increase in *PAX6* expression (fig. 1f). In addition, *OTX2, ZIC2, SOX9, HESX1, MSX2, DLX5, SOX4, SOX11*, and *SNAI2* were significantly activated during differentiation which demonstrates that the transcriptional program triggering progression towards NPCs was activated (Fig. 1f, Additional file 4: Figure S4h and Additional file 9: Figure S9a-c). Taken together, these results indicate that the induction cocktail effectively achieves efficient neural differentiation.

To measure the dynamic changes of c/s-regulatory elements at each differentiation stage, we performed ATAC-seq and chromatin accessibility analysis on bulk cells. These results showed widespread and comprehensive chromatin structure reprogramming during neural differentiation. In particular, TFBSs for several neural master regulators were enriched in temporally dynamic ATAC peaks, indicating that changes in chromatin accessibility are indeed associated with, and are probably responsive to, the regulation of neural-related TFs. In addition, we also investigated closing (lost) peaks dynamics as well as the functional annotation study, which was in line with the corresponding annotation of novel peaks (Additional file 2, 3: Figure S2, 3). We further identified several enriched TF motifs (e.g., *Pax2* in Ros-L and *FOXO1* in NPCs) (Additional file 17: Figure S17d, e) which are known to play an important role in neural differentiation, consistent with results from previous studies [39, 68].

By integrating single cell-based transcriptome profiling of 391 cells from five differentiation stages, we identified a variety of TFs that were differentially expressed throughout the differentiation process and showed distinct expression profiles among specific cell stages. The TFs *SOX2, PAX6, OTX2, SOX4, ZIC2, LHX5, HESX1*, and *SIX3* were significantly highly expressed at the EB stage (fig. 1f). It has been reported that members of the grainyhead-like (Grhl) family of TFs, which are well-conserved from *Drosophila* to human, are highly expressed during neurulation in mice and that a Grhl3-hypomorphic mutant resulted in NTDs [32, 67]. Remarkably, our results showed that two human Grhl family TFs, *GRHL2* and *GRHL3*, were significantly highly expressed at EB and Ros-E stage, respectively (Fig. 1f and Additional file 4: Figure S4h), and the downstream targets of *GRHL2* (including *E-CADHERIN*, also known as *CDH2)* were highly expressed at the neural rosette stage (fig. 1b) supporting a role for Grhl TFs in neural tube closure in humans. In addition, previous studies have shown that in the *Drosophila* olfactory system the homeobox gene *distal-less* is required for neuronal differentiation and neurite outgrowth [34]. Our data showed that four homologs of *distal-less (DLX3, DLX4, DLX5, DLX6)* were significantly up regulated at the Ros-E stage and were highly expressed in the Ros-E2 subpopulation (Fig. 1f and Fig. 2b) implying that the *distal-less* gene family plays a role in neural differentiation in humans.

We also applied scRNA-seq to our *in vitro* neural model to dissect the subpopulations present at each differentiation stage (Fig. 2 and Additional file 5-8: Figure S5-8). We were then able to reconstruct a differentiation trajectory based on the subpopulations that we identified by variable TF expression within each stage (fig. 3a). Strikingly, a divarication within the rosette stage across the differentiation trajectory was observed. Comparing Branch 1 to Branch 3, Branch 3 possessed the relatively lowly-expressed TFs *LHX5, HESX1* and *SIX3* (reported as anterior forebrain markers), as well as other crucial neural TFs (SOX2, *HMGB2, ZIC2, OTX1, FEZF1);* and the relatively highly-expressed TFs *TFAP2B, SOX9, ELK3*, and *SNAI2* (Fig. 3d, e and Additional file 9: Figure S9a, c) which are considered to be neural crest markers [53]. Though *SNAI2* was also expressed at the NPCs stage, combined with other neural crest markers, we proposed that Branch 3 was progressing more towards to neural crest cells (Fig. 3a-c and Additional file 9: Figure S9a-c). Taken together, these observations imply that the main differentiation trajectory (Branch 1 and Branch 2) is heading towards CNS, whereas Branch 3 is progressing towards neural crest cells.

It is important to note that the current scRNA-seq method by its nature only provides a snapshot of the gene expression profile for individual cells. A possible resolution for the above problem is to capture the sample with much more precise time points, which may, to some extent, overcome this limitation. Thus, in spite of the very interesting heterogeneity and cell fate commitment study inferred above, we cannot exclude the following factors that may affect cell subset identification in the above description; 1) temporal transcriptional states during transient differentiation process; 2) differentiation efficiency; and lagging and leading cells remaining in the differentiation process. However, we propose that the subsets dissection analysis facilitates a more precise description of the factors defining the differentiation trajectory. When we constructed the differentiation trajectory using the cells that collected at different time points, the results showed that all subpopulations in stages from iPSCs to NPCs followed a sequential differentiation process where each stage exhibited a relatively discriminative region with some of the subpopulations overlapping (fig. 3a), indicating that in spite of the above concerns, the trajectory was established by the natural features of the respective subsets and which is also supported by the observations that Ros-L2 possessing many early neural differentiation TFs, such as *SOX2, OTX2, PAX6, OTX1*, and *LHX5*, as well as forebrain markers (e.g., *HESX1)* and pluripotency-related TFs *(NANOG, SALL4, PRDM14)* (Additional file 7: Figure S7) were located in the reconstructed trajectory prior to the generation of Ros-E populations. In addition, we carried out the cell fate commitment analysis using Branch1, Branch2 and Branch3 which were grouped based on the cell locations on the trajectory rather than cell subsets identified by Seurat in order to minimize the above concerns.

Notably, our study reveals the regulatory network of TFs that are differentially expressed among neighbouring cell subpopulations to be likely candidates for promotion of cell fate transition. Based on the topology of this network, we focused on novel regulators *(PRDM1* and *ARID3A)*, especially *PRDM1*, which are located on the hub of the network, interacting with both known and novel neural regulators. Although the roles of several TFs have been reported during neural differentiation and brain pattering formation in humans, some TFs have been proposed to play a role in neural fate commitment in non-human species (mouse and zebrafish). However, the interaction partners, c/s-regulatory elements, and genetic regulatory networks of those TFs are yet to be resolved. Here, we identified the c/s-regulatory elements for *PRDM1* and *ARID3A* genes and predicted their upstream regulators. Of particular interest, *TFAP2C’s* role in regulating neural development has been widely reported, increasing the confidence of our predictions. In humans, *PRDM1* is reported to promote germ cell fate by suppressing neural effector *SOX2*, but the function of *PRDM1* in neural development is unknown. In zebrafish, *Prdm1a*, the homolog of the *PRDM1* gene, directly activates *foxd3* and *tfap2a* during neural crest specification [57]. Mutation of *prdm1* in zebrafish resulted in severe phenotypes with a decrease in the quantity of neural crest cells and the reduction in the size of structures derived from the neural crest [57]. Similarly, strong expression of *prdm1* was observed in the neural plate border of a basal vertebrate linage, lamprey, implying that the role of *prdm1* in the neural crest formation is likely a conserved, ancestral role [70]. Conversely, *prdm1* is dispensable for neural crest formation in mice, and instead is required for primordial germ cell specification suggesting that the neural crest specification function of *prdm1* in mice has been lost [71]. Overall, previous studies suggest that functions of *prdm1* are quite diverse and need to be investigated in species-, developmental-, and environmental-specific manners. Based on the known interaction between *PRDM1* and *SOX2* in humans, as well as the observation that *PRDM1* expression increased significantly from Ros-E2 to Ros-L3 and was preferentially expressed in Ros-L3 compared to other two subpopulations in the rosette stage (Fig. 4g, h; Additional file 7: Figure S7a, b and Additional file 10: Figure S10), we propose *PRDM1* as a novel neural regulator in early human neural differentiation. Our hypothesis is supported by the GO term and KEGG enrichment analysis of putative targets of *PRDM1*, which are significantly enriched in “axon guidance” and hippo pathway-associated terms (Fig. 4e and Additional file 13: Figure S13a). However, the functions of putative TFs need to be further investigated using experimental methods.

To infer cellular interactions, communication network analysis was applied to the expression profiles of ligands and receptors in stage-specific subpopulations. Two trends were observed in our cellular interaction network analysis: 1) the frequency of cellular interactions peaked at EB stage; and 2) different cell subpopulations showed a certain degree of specificity in their ligand-receptor spectrum. The observation that most interactions were inferred at the EB stage likely reflects the extensive cellular communication during embryogenesis and early neural differentiation (Additional file 14: Figure S14). Regarding the ligand-receptor expression spectra, matched ligand and receptor expression probably underlies the common functions shared by different cell subpopulations within the same stage. In contrast, those specific ligands or receptors probably reveal the unique regulatory code of distinct cell subpopulations. For example, *WNT5A*, a crucial regulator of neurogenesis during the development of cerebellum, and *BMP4*, one of the key regulators of dorsal cell identity in the neural tube [72], were highly expressed in Ros-L1 compared to other cell subpopulations (Fig. 5c). FZD5 (required for eye and retina development in mouse [73]), and *FGF19* (required for forebrain development in zebrafish [74]) were preferentially expressed in Ros-L2 (Fig. 5d and Additional file 22: Table S4). *WNT7A*, involved in several aspects of neurogenesis, including synapse formation and axon guidance [75] and *FGF1*, which maintains the self-renewal and proliferation of NPCs [76], were specifically expressed in Ros-L3 (Additional file 22: Table S4). Pavličev et al. inferred the cell communication network of the maternal-fetal interface and found that ligand-receptor profiles could be a reliable tool for cell type identification [65]. Consistent with their findings, our study suggests that the repertoire of ligands-receptors in neural cell types could probably, to some extent, represent the identity of cell subpopulations.

There might be a concern that we only used one genetic background cell line for this study, possibly making the cogency of our findings limited. To address this, we performed ESCs neural differentiation and captured bulk transcriptome profiles of the corresponding differentiation stages (ESCs, EB, Ros-E, Ros-L and NPCs). The observations in ESCs were reproducible in iPSCs with regards to 1) PCA analysis (Additional file 18: Figure S18a); 2) with a high Pearson correlation coefficient between the corresponding cell stage derived from iPSCs and ESCs (Additional file 18: Figure S18b); and 3) validation analysis of subset-specific markers (MAFB, SOX9, PRDM1 and NR2F1). In addition, novel neural TF (PRDM1) expression in different genetic cell lines (H1_ESCs, H7_ESCs, H9_ESCs, iPS25 and iPS129) was consistent with the above heterogeneity study (Additional file 18: Figure S18c, d, e). Together, our findings are supported by different genetic cell lines mitigating the concern that our results are limited to the cells forming the basis of this study.

Through differential expression analysis, we identified genes specifically expressed at each stage which include both cell status master regulators such as TFs and signalling components, as well as realizators [24] which could directly determine cell growth, cell proliferation, cell morphology and cell-cell interaction. Within each stage, we identified subpopulations with distinct expression signatures, which might represent functional cell clusters or transient cell state given that neural cells have been shown to demonstrate significant heterogeneity as they express different surface proteins, exhibit diversified morphologies and secrete a variety of cytokines. Therefore, it is necessary to explore the heterogeneity of cell subpopulations and study each subpopulation in a case-by-case manner. In summary, our data show conclusively that both transcriptome and regulome dramatically change during neural differentiation, which affects a variety of biological pathways crucial for neural differentiation. We also propose several putative TFs as well as the ligands-receptors interaction spectrum that are important in each differentiation stage which paves the way for a deeper understanding of the cell fate decision and regulatory mechanisms driving the differentiation of the neural lineage.

## Materials and methods

### Ethics statement

The study was approved by the Institutional Review Boards on Ethics Committee of BGI (Permit No.BGI-IRB 14057). The participant (dermal fibroblast, Fib129) signed informed consent and voluntarily donated the samples for our study.

### Cell culture and reprogramming

The human fibroblast cell line was derived from the dermal skin of a healthy female donor with written informed consent. Briefly, the skin tissue was washed with DPBS several times, sliced into approximately 1mm or smaller fragment size, enzymatically dissociated in High Dulbecco’s modified Eagle medium (H-DMEM, Gibco, 11965118) with 100U/ml collagenase type IV incubating in 37°C overnight, then 0.05% trypsin incubating for 5 min. The dissociation was terminated by adding 2 ml fibroblast cell culture medium (H-DMEM +10% FBS + 5ng/ml bFGF+ 2mM Gln) followed by centrifugation at 300g for 5 min. The cells were resuspended with fibroblast cell culture medium, and cultured at 37°C in a 5% CO_2_ incubator. The fibroblast cell culture medium was changed every 2 days until reaching 80%-90% confluence and cells were passaged every 3-4 days.

For reprogramming, non-integrative human iPSCs were generated following a modified Shinya Yamanaka method [77]. Briefly, 5×10^5^ human fibroblast cells at passage 4 were nucleofected with the program for human dermal fibroblast NHDF (Lonza, CC-2511) with 2.4ug episomal plasmids, including pCXLE-hOCT3/4-shp53-F (Addgene, 27077), pCXLE-hSK (Addgene, 27078), pCXLE-hUL (Addgene, 27080). Transfected cells were cultured in a six-well plate with culture medium containing H-DMEM supplemented with 10% FBS. The cells were trypsinized and 1×10^5^ cells were seeded onto a 10cm^2^ dish covered with feeder and cultured in a medium containing H-DMEM with 10% FBS while reaching 80% confluence. After that, the medium was changed to hiPSCs medium containing DMEM/F12 (Gibco, 11320-033), 20% KSR (Gibco,10828-028), 2mM L-glutamine (Sigma, G8540), 0.1jM NEAA (Gibco,11140-050), 0.1|jM P-Mercaptoethanol (Gibco, 21985-023) and 10ng/ml human bFGF (Invitrogen, PHG0021). The iPSCs colonies were picked at around day 25 and maintained in hiPSCs medium.

### Neural differentiation

We applied a well-adopted neural differentiation protocol [8,16]. Briefly, human iPSCs were maintained as described above. To induce neural rosettes, hiPSCs were mechanically picked and washed with DMEM/F12 twice, and then cultured for 4 days in suspension with 5μM dorsomorphin (Sigma, P5499) and 5μM SB431542 (Sigma, S4317) in hiPSCs medium without bFGF for embryoid bodies (EBs) formation, then the EBs were attached on matrigel (BD, 354277) coated dishes (BD, 354277) and cultured in DMEM/F12 (Gibco, 11320-033) supplemented with 20 ng/ml bFGF, 1*N2 (Gibco, 17502-048) and 2ug/ml heparin (Sigma, 1304005) for an additional 3 or 5 days to harvest rosette-early (Ros-E) and rosette-late (Ros-L) cells, respectively. To collect neural progenitor cells (NPCs), rosettes structure that appeared in the center of attached colonies at Ros-L stage were carefully harvested using pulled glass pipettes and seeded on matrigel-coated dishes and cultured in DMEM/F12 supplemented with 1x N2, 1x B27 (Gibco,12587-010), 20 ng/ml bFGF, 20 ng/ml EGF (Invitrogen, PHG0311) and 2ug/ml heparin (Sigma,1304005) for additional 7 days, and the medium was changed every 2 days. At day 16, the NPCs reaching approximately 80% confluence were collected, and all the mass or adherent cell samples were treated with TrypLE™ Express Enzyme (Gibco, 12604-021) for single cell dissociation and cryopreservation in gas-phase liquid nitrogen for further sequencing.

### Immunofluorescence staining

HiPSCs and Ros-L cells were fixed in 4% paraformaldehyde in DPBS for 20 min and permeabilized with 1% Triton X-100 for 20 min at room temperature. After 60 min blocking with 2% normal goat serum, hiPSCs were incubated with primary antibodies OCT4 (1: 200, Abcam), NANOG (1: 200, Abcam), and Ros-L cells were incubated with primary antibodies PAX6 (1: 200, Abcam), SOX2 (1:200, Abcam), NESTIN (1: 200, Abcam), SOX1 (1: 200, Abcam), Zo-1 (1:100, Abcam) and N-CAD (1: 100, Abcam) overnight at 4 °C, then stained with secondary antibodies (goat anti rabbit IgG-Cy3 diluted1: 300 and goat anti mouse IgG-Cy3 diluted 1: 300) for 60 min at room temperature. DAPI (1: 500) was used as counter-staining for nuclei. The images were captured and analyzed with the Olympus IX73 and Image J.

### Single cell RNA sequencing

Cells at indicated time points were collected for single cell RNA-seq and global transcriptome analysis. TrypLE™ Express Enzyme (Gibco, 12604-021) was applied for single cell dissociation. Single-cell RNA-seq library construction was conducted according to an automated pipeline called microwell full-length mRNA amplification and library construction system (MIRALCS) as described previously [78]. 50bp single-end sequencing was performed using the BGISEQ-500 platform.

### Assay for transposase-accessible chromatin sequencing (ATAC-seq)

We profiled open chromatin accessibility sequencing (ATAC-seq) of neural differentiation process for five stages including iPSCs, EB, Ros-E, Ros-L and NPCs samples. ATAC-seq libraries were prepared using a modified protocol based on previous study [79]. Briefly, 50,000 cells were collected for each sample, washed with pre-cooling PBS and resuspended in 50 jl of ice-cold lysis buffer (10 mM Tris-HCl, pH 7.5, 10 mM NaCl, 3 mM MgCl2, 0.1% IGEPAL CA-630). Permeabilized cells were resuspended in 50 jl transposase reaction buffer (1* TAG buffer, 2.0 jl Tn5 transposes enzyme) and incubated for 30 min at 37 °C. PCR amplification and size selection (150-500 bp) were performed using Agincourt AMPure XP (Beckman Coulter) and Bioanalyzer 2100 (Agilent). Libraries were pooled at equimolar ratios with barcodes and sequenced on BGISEQ-500 platform.

### Pre-processing and quality control of single cell RNA-seq

The original FASTQ data of the 527 samples were aligned to the rRNA database (downloaded from NCBI) to remove rRNAs and the remaining reads were processed with SOAPnuke (version 1.5.3) [80] to trim adaptors and filter out the low-quality reads. The filtered data were aligned to the reference genome (hg19) using hisat2 (HISAT2 version 2.0.1-beta) [81]. Reads were counted using the R package GenomicAlignments [82] (mode=‘Union’, inter.feature=FALSE), and normalized to RPKM with edgeR [83]. Cells were filtered using following parameters: genome mapping rate more than 70%, fraction of reads mapped to mitochondrial genes less than 20%, mRNA mapping rate more than 80%, ERCC ratio less than 10%, and gene number more than 5000. Further, correlation of ERCC among cells was used to evaluate the quality of each cell (threshold=0.9). At last, 445 single cells remained for further analysis in this project.

### Identification of differentially expressed genes

Differential expression of genes in iPSCs (n = 71 cells), EB (n = 57 cells), Ros-E (n = 81 cells), Ros-L (n = 92 cells), and NPCs (n = 90 cells) was determined using SCDE (single cell differential expression analysis) [84] with default parameters except requiring a minimum of 100 genes (parameter min.lib.size = 100 to call scde.error. models function). The Z scores and corrected Z scores (cZ) to adjust for the multiple testing were converted into two-tailed p-values and adjusted to control for FDR using pnorm function in R. The significantly differentially expressed genes were selected based on following criteria: adjusted p-value < 0.01 and fold-change > 2.

### Constructing trajectory using differentially expressed genes

Monocle [85] ordering was conducted for all iPSCs, EB, Ros-E, Ros-L and NPCs cells using the set of variable genes with default parameters except we specified reduction_method =“DDRTree” in the reduceDimension function. The variable genes were selected using the Seurat R package [86].

### Analysis of heterogeneity in each cell stage

The heterogeneity of each cell stage was determined using Seurat R package [86] by the normalized expression level of reported transcription factors (retrieved from AnimalTFDB 2.0) [89]. Briefly, PCs with a p-value less than 0.01 were used for cell clustering with reduction.type=“pca” and resolution=“1.0”. The FindallMarkers function of Seurat package was used to identify marker genes for each cluster using default parameters.

### ATAC peak calling

We aligned ATAC-seq data to hg19 using Bowtie2 [88] and called peaks using MACS2 [89]. We established a standard peak set by merging all overlapping peaks. The IDR pipeline [90] was used to identify reproducible peaks between two biological replicates. Only peaks with IDR<=0.05 were considered reproducible and retained for downstream analysis. Pearson correlation coefficients of two biological replicates at each stage were calculated. Stage-specific peaks were defined as peaks having no overlap with any peaks in other stages. Novel peaks were defined as peaks non-overlapping with previous stages. In the case of iPSCs, all peaks were annotated as novel peaks.

### Targets assignment of ATAC peaks

For reproducible peaks, we applied HOMER [91] to assign putative targets for peaks. For stage-specific peaks, ChIPseeker [92] was used for putative target assignment. In both strategies, the putative target of a certain peak is defined as the gene with TSS closest to the peak summit location.

### GO term and KEGG enrichment analysis

Lists of genes were analysed using DAVID [93,94] and the BH method was used for multiple test correction. GO terms with a FDR less than 0.01 or 0.05 were considered as significantly enriched. Target genes of stage-specific ATAC peaks were analysed using the R package, clusterProfiler [95], in which an adjusted p-value of 0.05 was used to identify significantly enriched GO and KEGG terms associated with each set of peaks.

### Regulatory network construction

The scRNA-seq profiles among each cell types were compared using SCDE package [84]. TFs significantly differentially expressed, with adjusted p-value threshold of 0.05, among neighboring cell types were submitted to STRING database [64] to infer regulatory networks based on known interaction relationships (supported by data from curated databases, experiments and text-mining). TFs without any interactions with other proteins were removed from the network. To select key regulators, we used a threshold of 5 and all TFs with number of interactions above the threshold were considered as key regulators.

### Putative targets prediction, GO term and KEGG enrichment analysis

The target prediction and enrichment analyses were performed using the FIMO [96] and GREAT [97] packages, respectively. Briefly, the peak files in a certain stage were scanned for the presence or absence of TF motifs, which were downloaded from the Jasper database [98]. Genes with a TSS closest to TF motif-containing peaks were considered as putative targets of certain TFs.

### Construction of cellular communication network

The ligand-receptor interaction relationships were downloaded from the database, IUPHAR/BPS Guide to PHARMACOLOGY [98], and the Database of Ligand-Receptor Partners (DLRP) [65, 100]. The average expression level of TPM of 1 was used as a threshold. Ligands and receptors above the threshold were considered as expressed in the corresponding cluster. Adjusted *P* value of 0.05 was used as a threshold to identify ligands/receptors specifically expressed in a subpopulation. The R package Circlize [101] was used to visualize the interactions.

### Motif enrichment analysis

Motifs enriched in each set of ATAC peaks were identified using findMotifsGenome.pl from HOMER [91] using following parameters: -size −100,100 -len 4,5,6,7,8,9,10,11,12.

## Additional files

**Additional file 1: figure S1.**
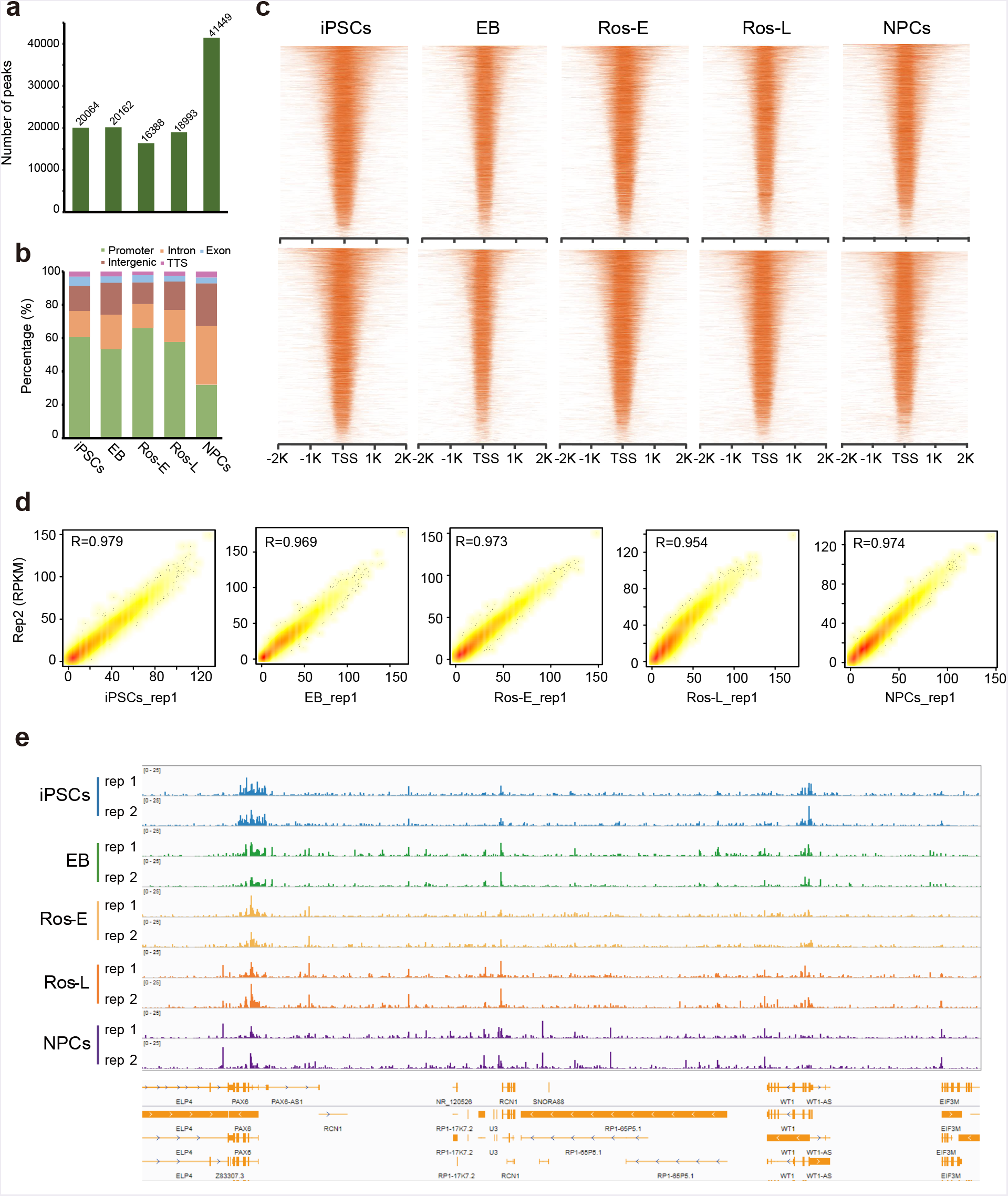
Quality control of ATAC-seq. **a** Bar graphs indicate the number of chromatin open regions detected at each cell stage of neural differentiation. **b** Genomic components (distribution) of the peaks in each cell stage during neural differentiation. **c** Heatmaps reporting the chromatin accessibility density within ±2 kb of TSSs. **d** Biological replicates of bulk ATAC-seq show high reproducibility. **e** IGV screenshot showing highly correlated ATAC signals in selected region between replicates.

**Additional file 2: figure S2.**
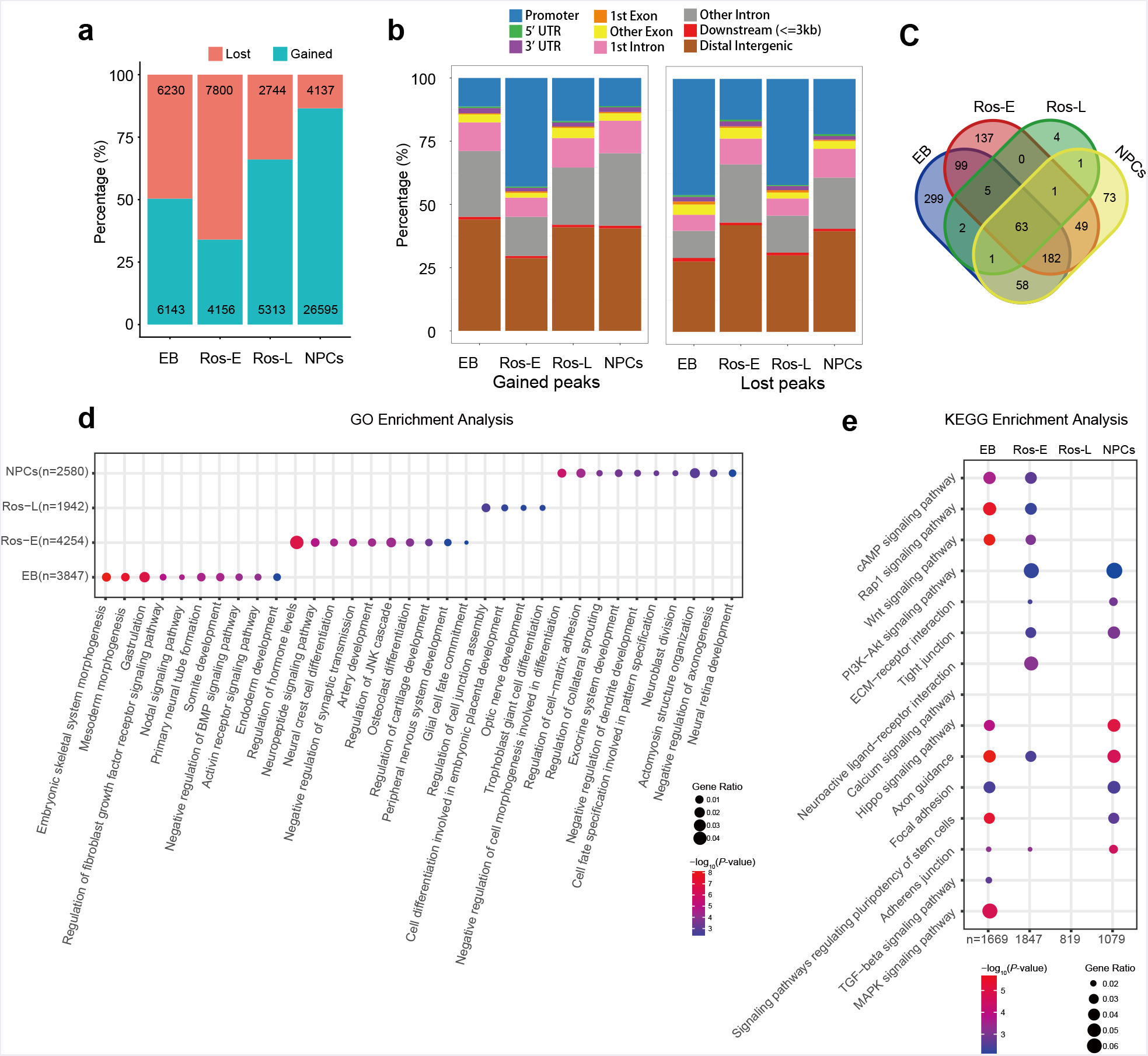
Dynamics of gained and lost peaks during neural differentiation. **a** Bar graph shows the number of gained and lost peaks at each cell stage. **b** Bar graph shows genomic composition of gained and lost peaks at each cell stage respectively. **c** Venn plot of GO enrichment analysis on the genes associated with lost peaks at each stage (adjusted *P*-value ≤ 0.01). **d** Selected GO terms identified by genes associated with lost peaks specific to the respective indicated cell stage (adjusted *P*-value ≤ 0.01). **e** Selected differential pathways identified by genes associated with lost peaks at indicated cell stages (adjusted P-value ≤ 0.01).

**Additional file 3: figure S3.**
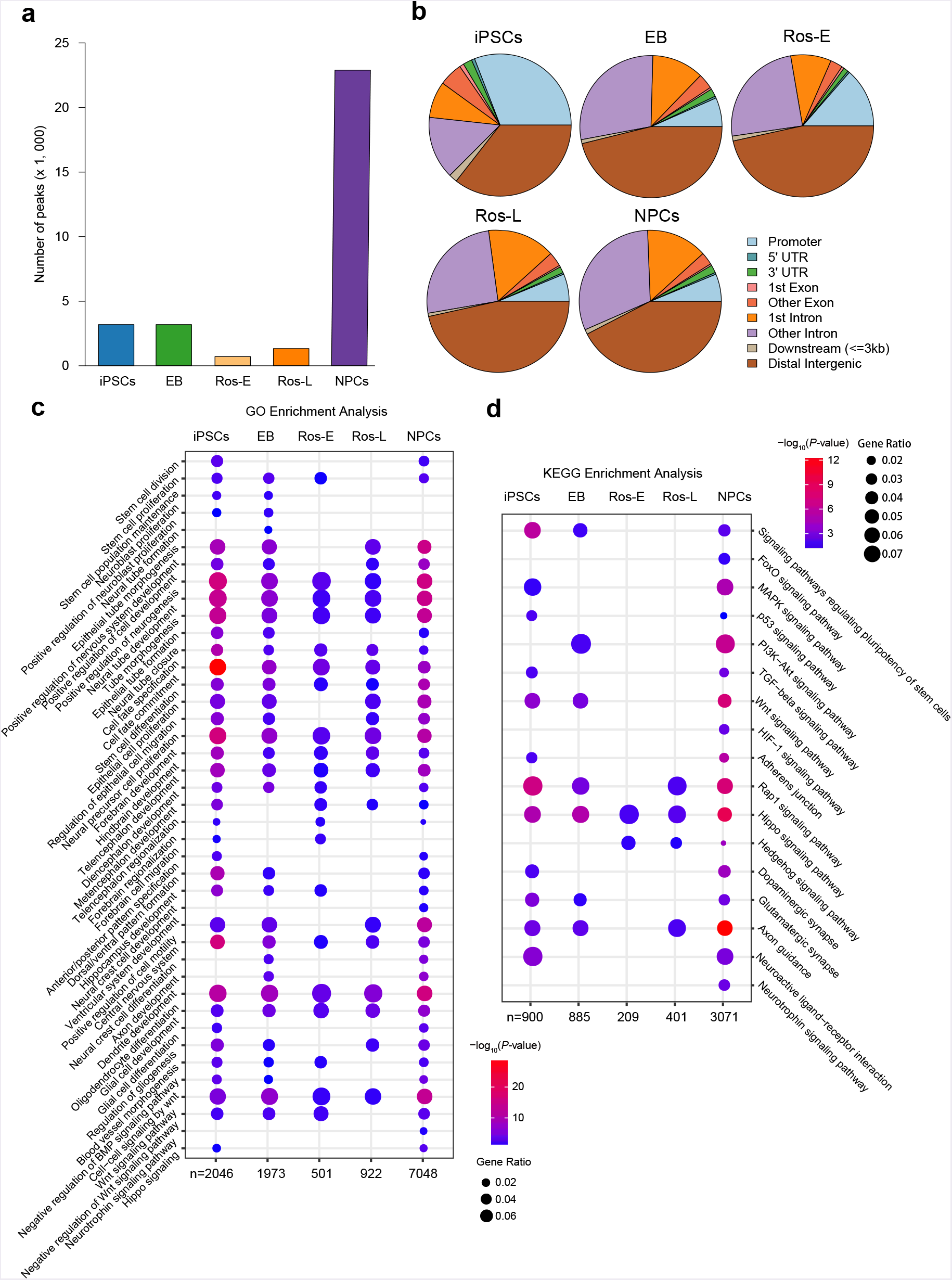
Stage-specific features of c/s-regulatory elements during neural differentiation. **a** Bar plot showing the number of stage specific ATAC peaks at iPSCs, EB, Ros-E, Ros-L and NPCs stage (adjusted *P*-value ≤ 0.01). **b** Pie chart shows genomic composition of stage specific peaks respectively. **c, d** GO term and KEGG enrichment analysis of stage specific peaks, respectively (adjusted *P*-value ≤ 0.05).

**Additional file 4: figure S4.**
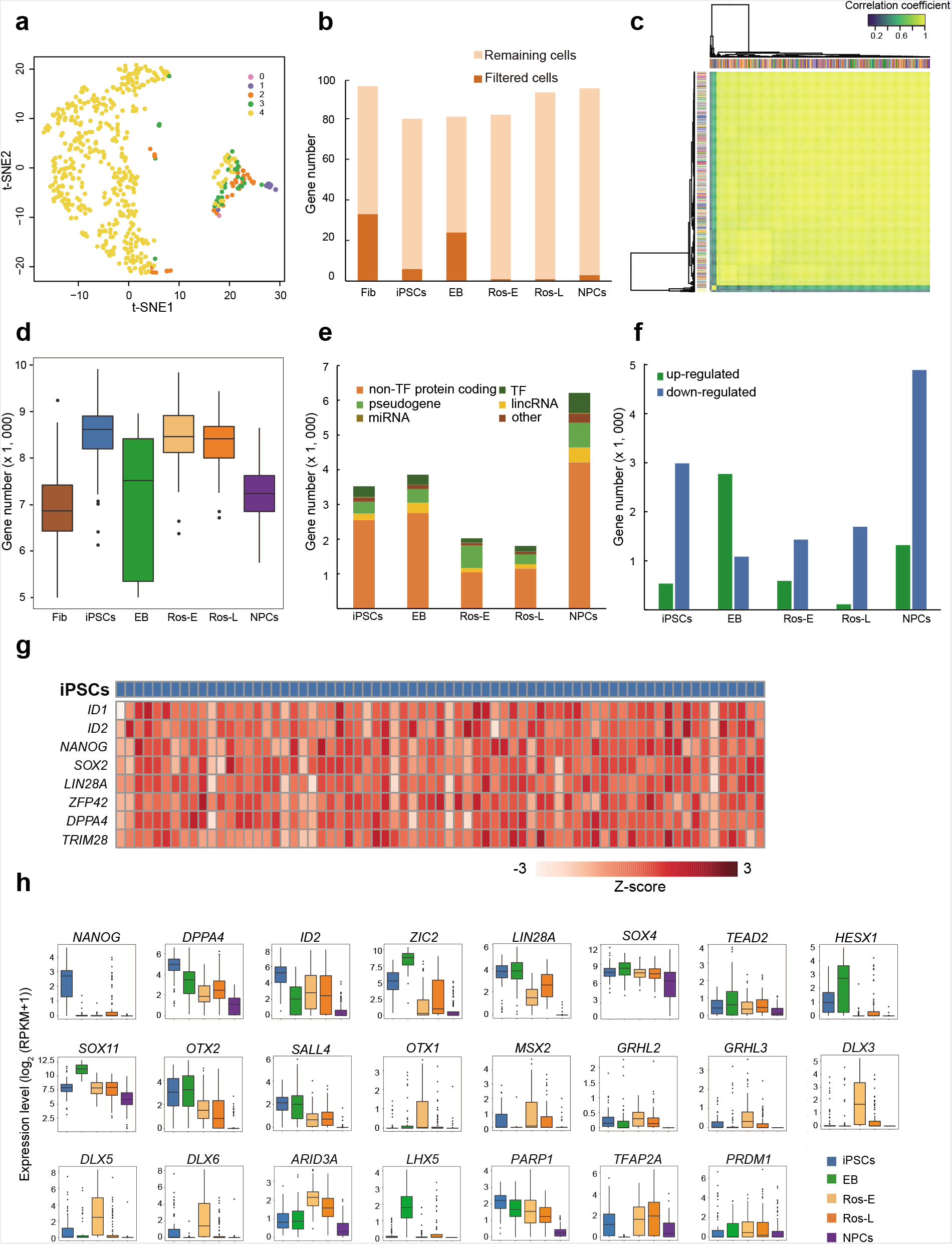
Quality control of scRNA-seq. **a** Graph indicates data quality of totally 527 single cells. Color scheme indicates the filter conditions, each dot represents one cell, and yellow dots showing the cells that successfully passed all criteria were used for downstream analysis. **b** Bar plots show the percentage of filtered cells and remaining cells. **c** ERCC correlation analysis of all single cells showing very little batch effects. **d** Box plots report the number of expressed genes for each cell stage after quality control filtering. Each dot represents an outlier gene and each box represents the median and first and third quartiles. **e** Genomic distribution of genes at each cell stage. **f** Summary of up-regulated and down-regulated genes at each cell stage compared to other stages. **g** Expression pattern of pluripotency-associated genes in iPSCs. Color scheme is based on z-score distribution from −3 (light red) to 3 (red). **h** Expression pattern of representative differentially expressed TFs during neural differentiation (adjusted *P*-value ≤ 0.01).

**Additional file 5: figure S5.**
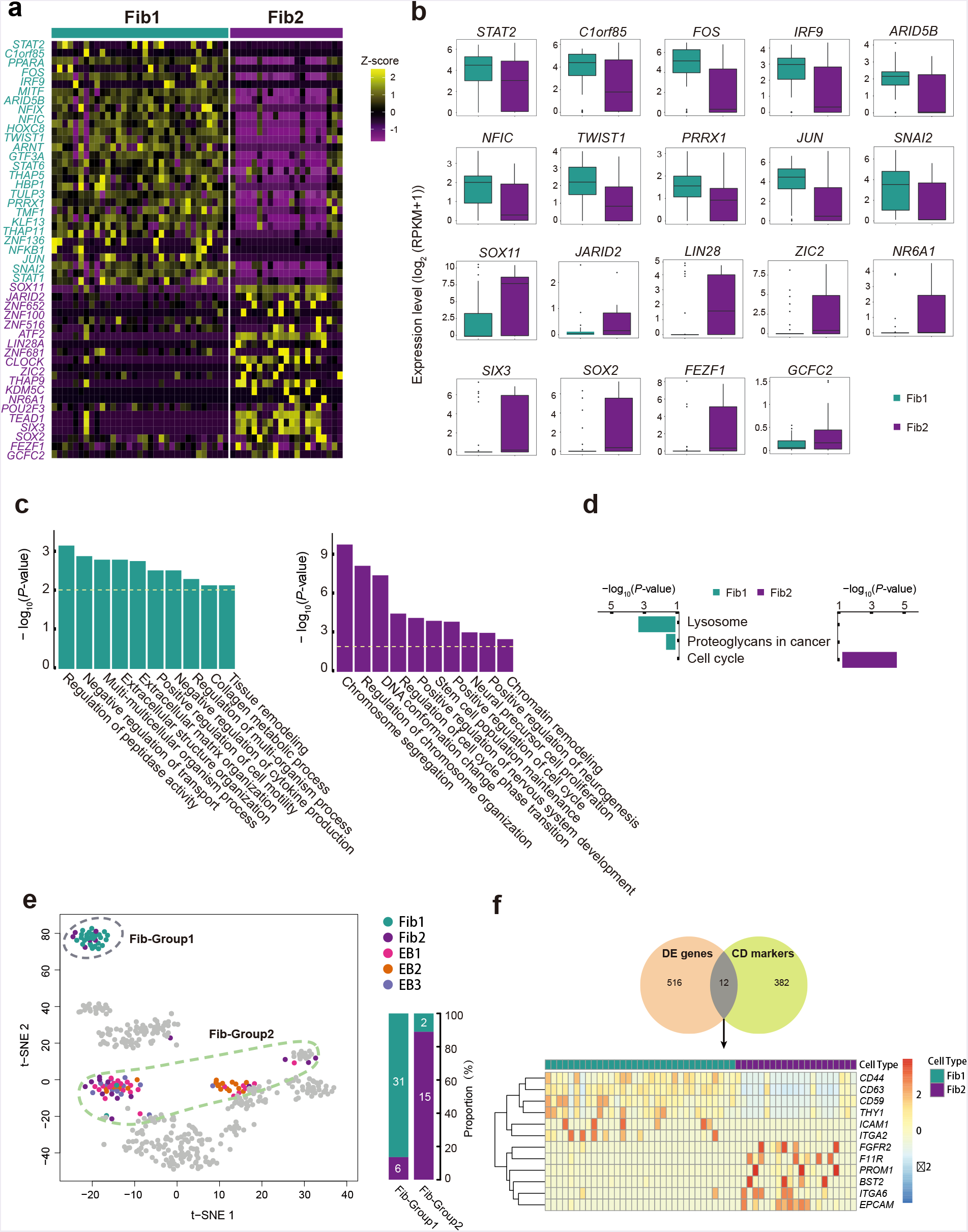
Subgroups identification and key transcriptomic features within Fib stage. **a** Heatmap reports scaled expression [log_2_ (RPKM+1)] of discriminative TF sets for each cluster in Fib stage with *P*-value cutoff ≤ 0.01. Color scheme is based on z-score distribution from −1 (purple) to 2 (yellow). Gene symbols highlight with color specific to the respective Fib subset. **b** Box plots of selected TFs defined in Figure S5a. **c** Selected GO terms identified by up-regulated genes specific to the respective Fib subpopulation with the color as indicated (Green: GO terms specific to Fib1; purple: GO terms specific to Fib2). **d** KEGG enrichment analysis of all terms in Fib subpopulation, respectively. **e** Fib-Group1 and Fib-Group2 based on their location on the t-SNE are marked by dashed ellipse. The columns represent the components of Fib-Group1 and Fib-Group2, respectively. **f** Comparison of differentially expressed (DE) genes between Fib subpopulation with CD markers dataset (HUGO Gene Nomenclature Committee, HGNC) and the heatmap of differentially expressed CD markers between the two Fib subpopulation.

**Additional file 6: figure S6.**
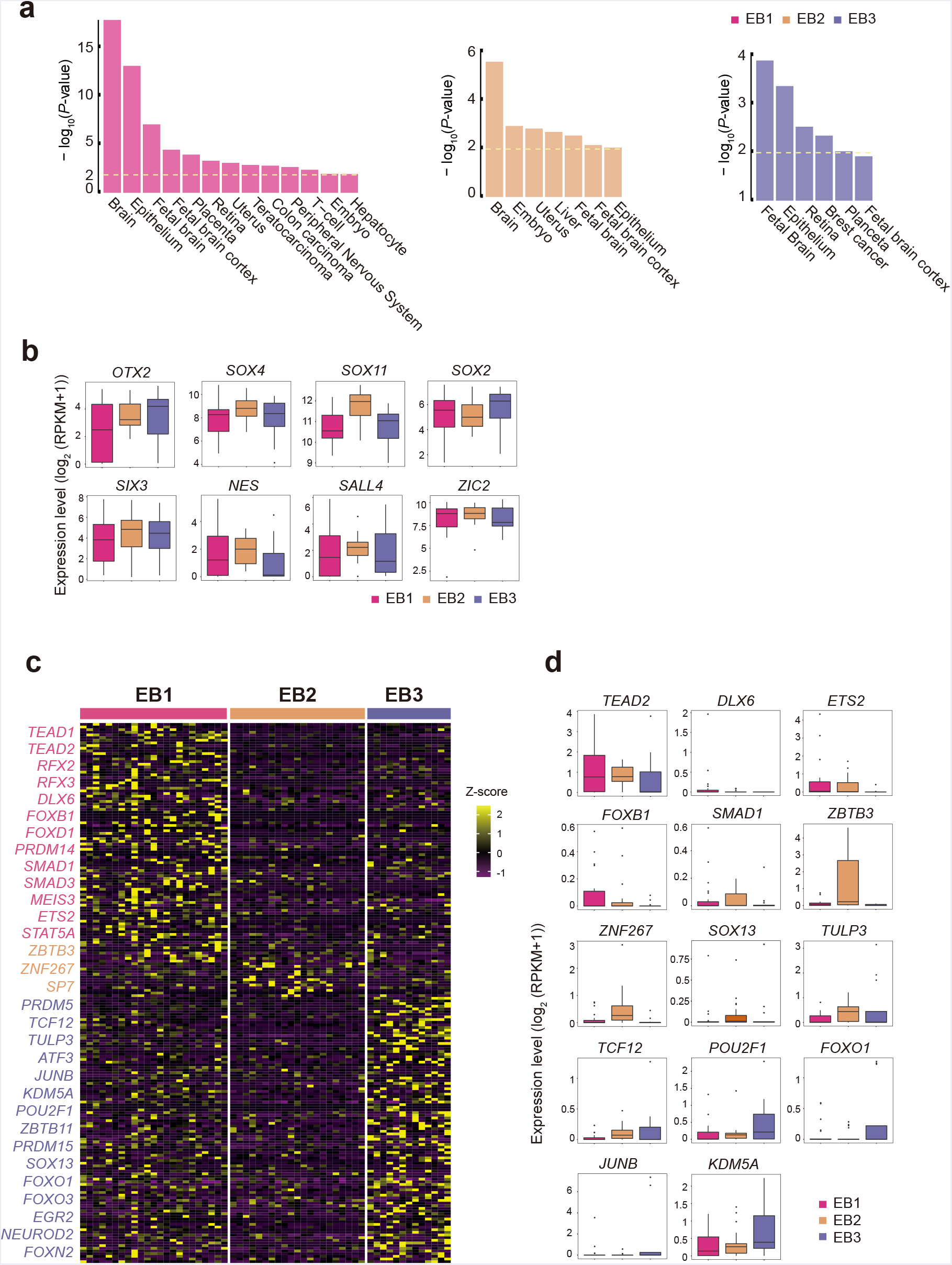
Subgroups identification and key transcriptomic features within EB stage. **a** David for tissue enrichment analysis of up-regulated genes defined by three EB subgroups compared to iPSCs stage respectively. **b** Box plots of commonly expressed genes across EB subsets. **c** Heatmap reports scaled expression [log_2_ (RPKM+1)] of discriminative TF sets for each cluster in EB stage with P-value cutoff ≤ 0.01. Color scheme is based on z-score distribution from −1(purple) to 2 (yellow). Gene symbols highlight with color specific to the respective EB subset. **d** Box plot of selected TFs defined in Figure S6a.

**Additional file 7: figure S7.**
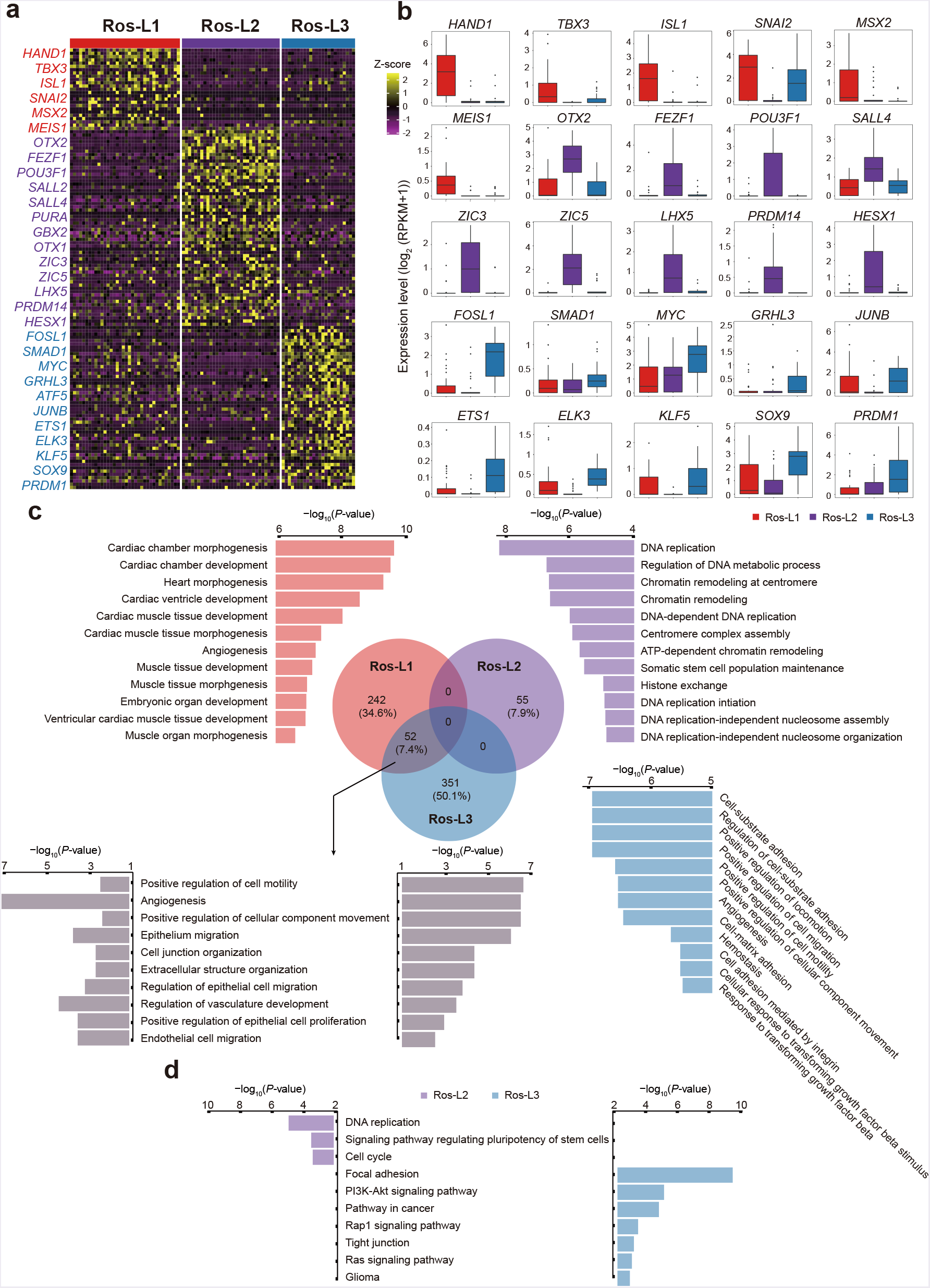
Subgroups identification and key transcriptomic features within Ros-L stage. **a** Heatmap reports scaled expression [log_2_ (RPKM+1)] of discriminative TF sets for each cluster in Ros-L stage with P-value cutoff ≤ 0.01. Color scheme is based on z-score distribution from −2 (purple) to 2 (yellow). Gene symbols highlight with color specific to the respective Ros-L subset. **b** Box plots of selected TFs defined in Figure S7a. **c** Top 12 of GO terms identified by up-regulated genes specific to the respective Ros-L subpopulation with the color as indicated (red: GO terms specific to Ros-L1; purple: GO terms specific to Ros-L2; blue: GO terms specific to Ros-L3; gray: selected GO terms shared by Ros-L1 and Ros-L3). **d** KEGG enrichment analysis of Ros-L2 (all terms) and Ros-L3 (selected terms), respectively.

**Additional file 8: figure S8.**
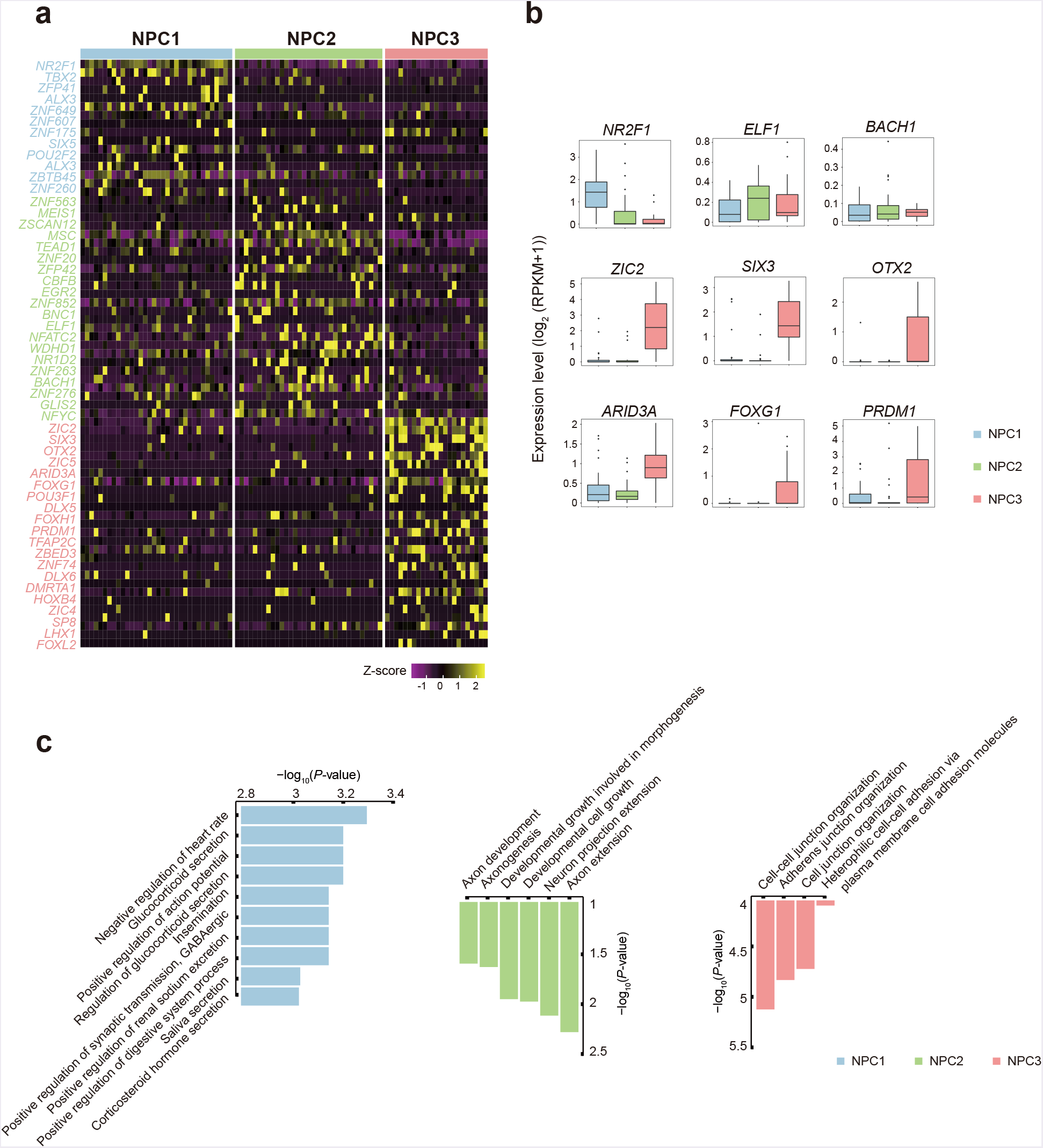
Subgroups identification and key transcriptomic features within NPCs stage. **a** Heatmap reports scaled expression [log_2_ (RPKM+1)] of discriminative TF sets for each cluster in NPCs stage with P-value cutoff ≤ 0.01. Color scheme is based on z-score distribution from −1 (purple) to 2 (yellow). Gene symbols highlight with color specific to the respective NPC subset. **b** Box plot of selected TFs defined in Figure S8a. **c** Top 10 (NPC1) and all (NPC2 and NPC3) of GO terms identified by up-regulated genes specific to the respective Ros-L subpopulation with the color as indicated (blue: GO terms specific to NPC1; green: GO terms specific to NPC2; pink: GO terms specific to NPC3).

**Additional file 9: figure S9.**
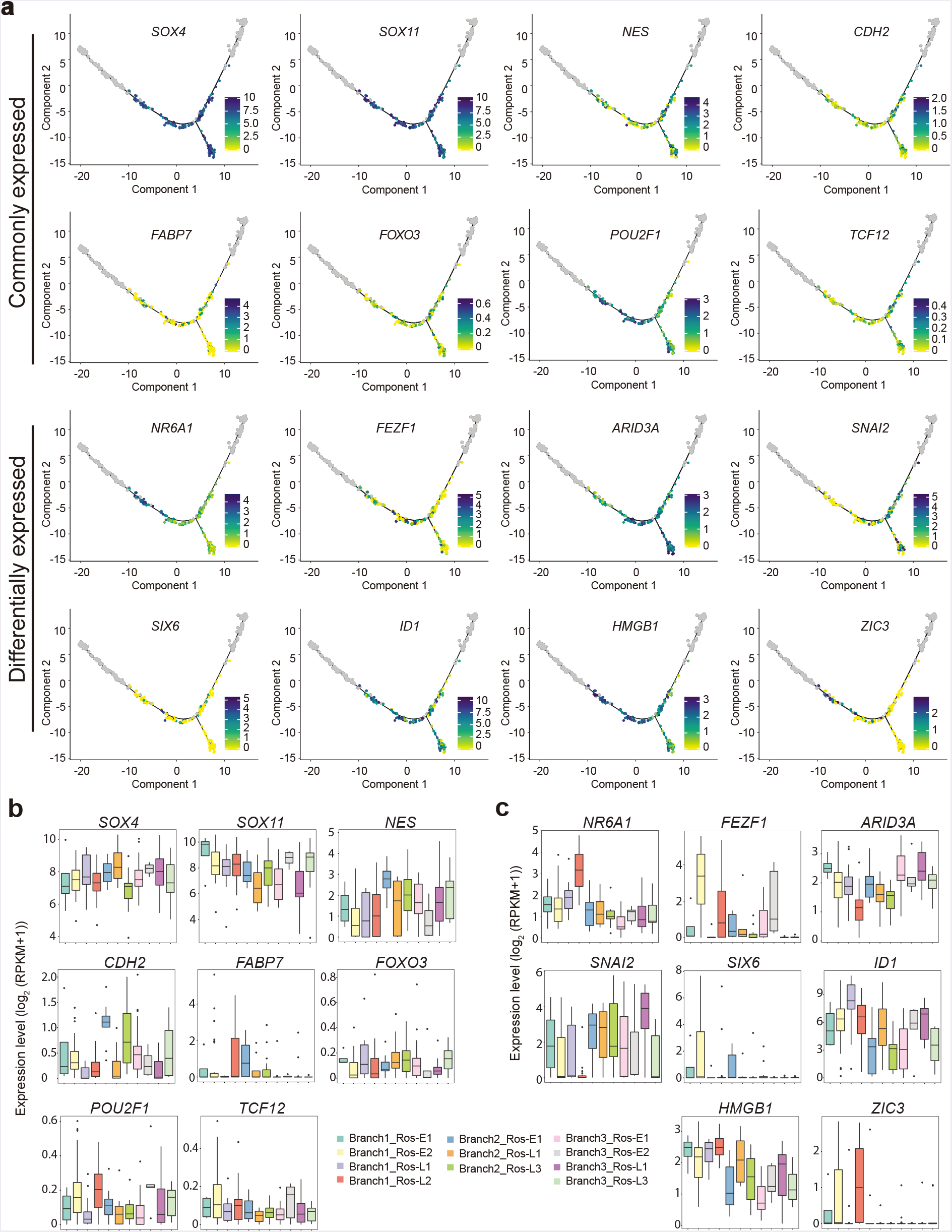
Expression pattern of selected transcription factors (TFs) within rosettes (Ros-E and Ros-L) stage. **a** Expression enrichment of commonly and differentially expressed TFs along the differentiation trajectory. Color scheme is based on expression [log_2_ (RPKM+1)]. **b, c** Expression pattern of selected TFs with respect to Figure S9a (adjusted *P*-value ≤ 0.01).

**Additional file 10: figure S10.**
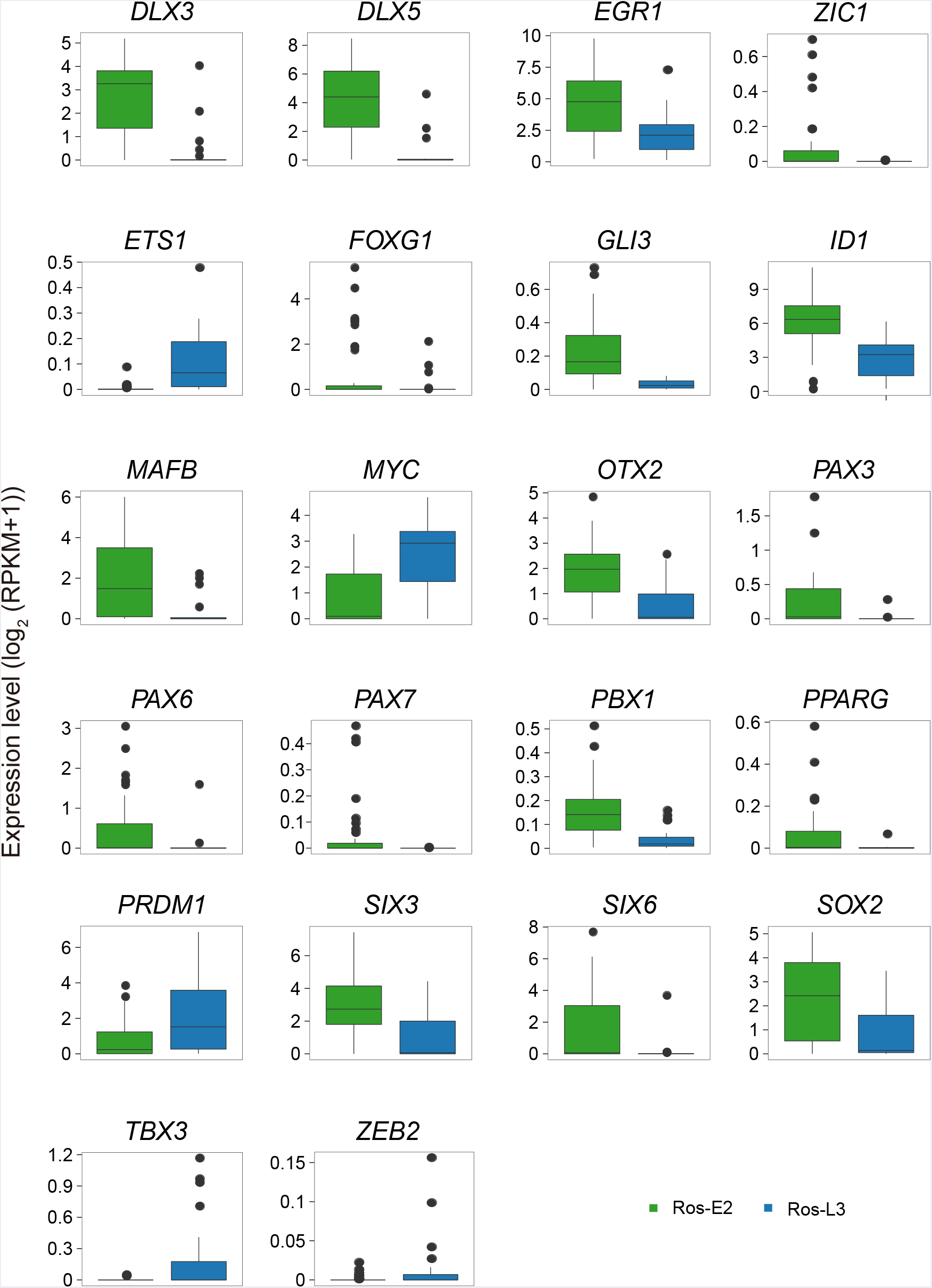
Differentially expressed transcription factors (TFs) between Ros-E2 and Ros-L3. Ros-E2 and Ros-L3 were shown in green and blue column, respectively (adjusted *P*-value ≤ 0.01).

**Additional file 11: figure S11.**
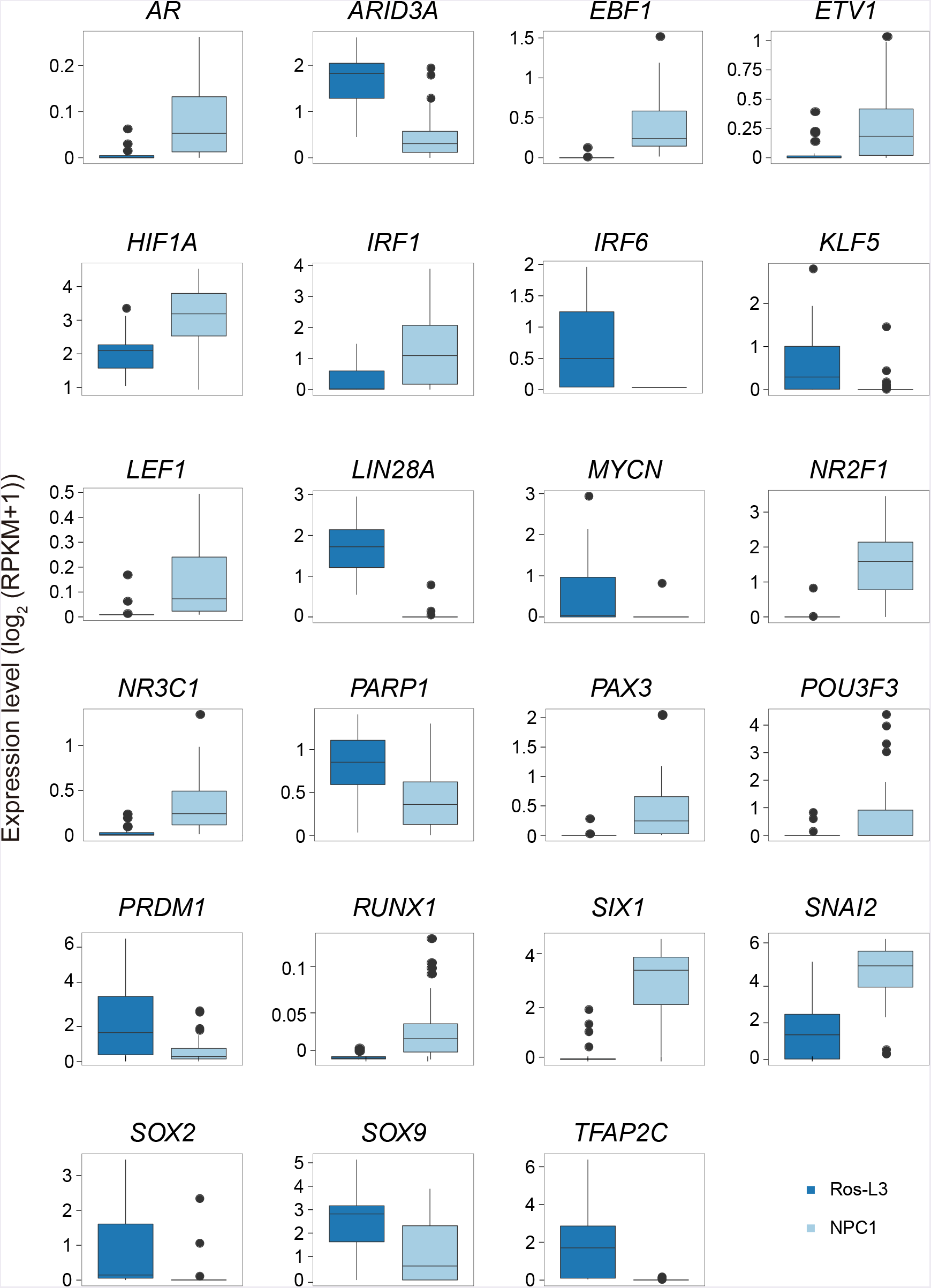
Differentially expressed transcription factors (TFs) between Ros-L3 and NPC1. Ros-L3 and NPC1 were shown in dark blue and light blue column, respectively (adjusted *P*-value ≤ 0.01).

**Additional file 12: figure S12.**
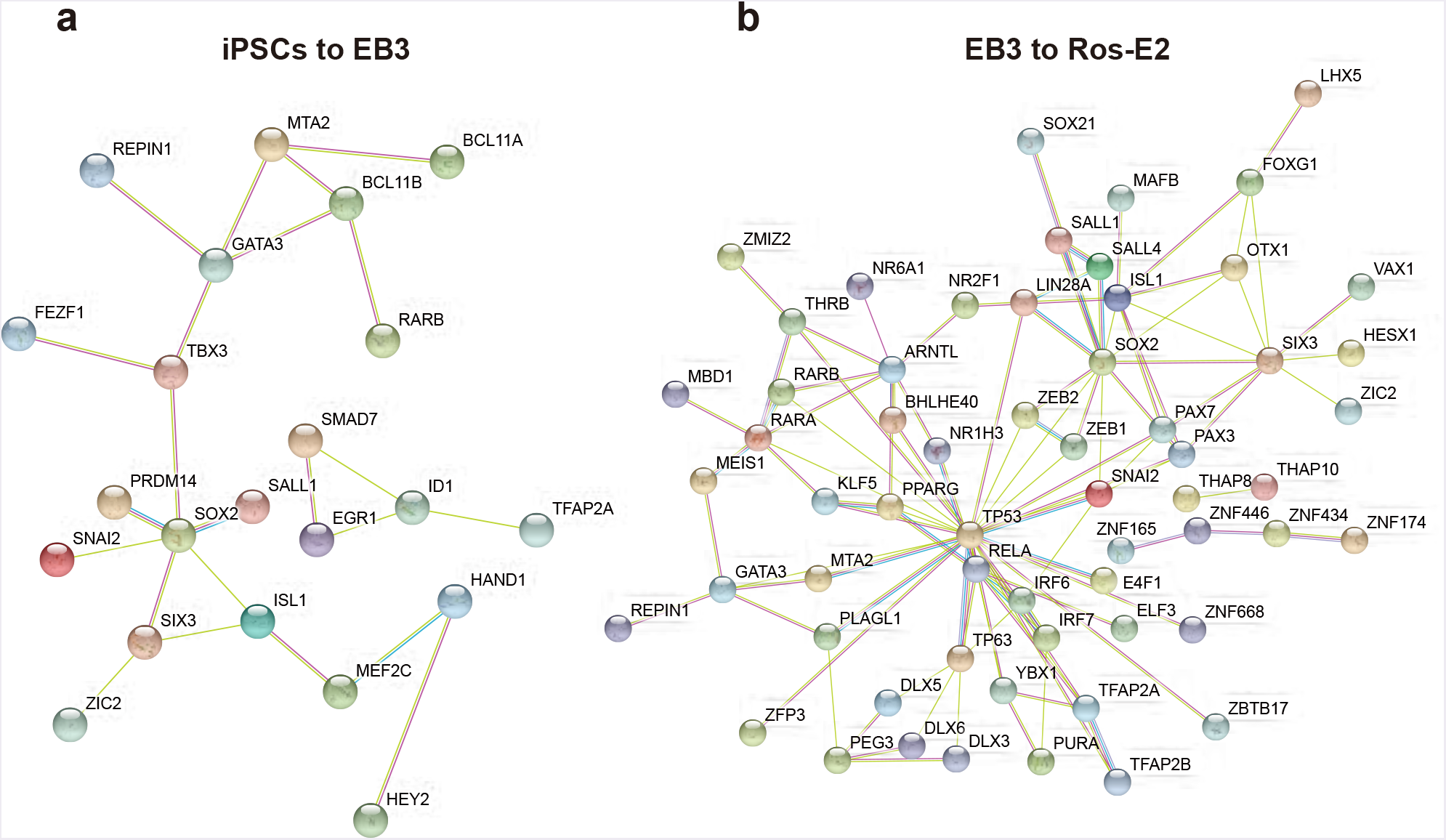
Key regulators during neural differentiation. **a** Regulatory network of differentially expressed TFs between iPSCs and EB3. **b** Regulatory network of differentially expressed TFs between EB3 and Ros-E2.

**Additional file 13: figure S13.**
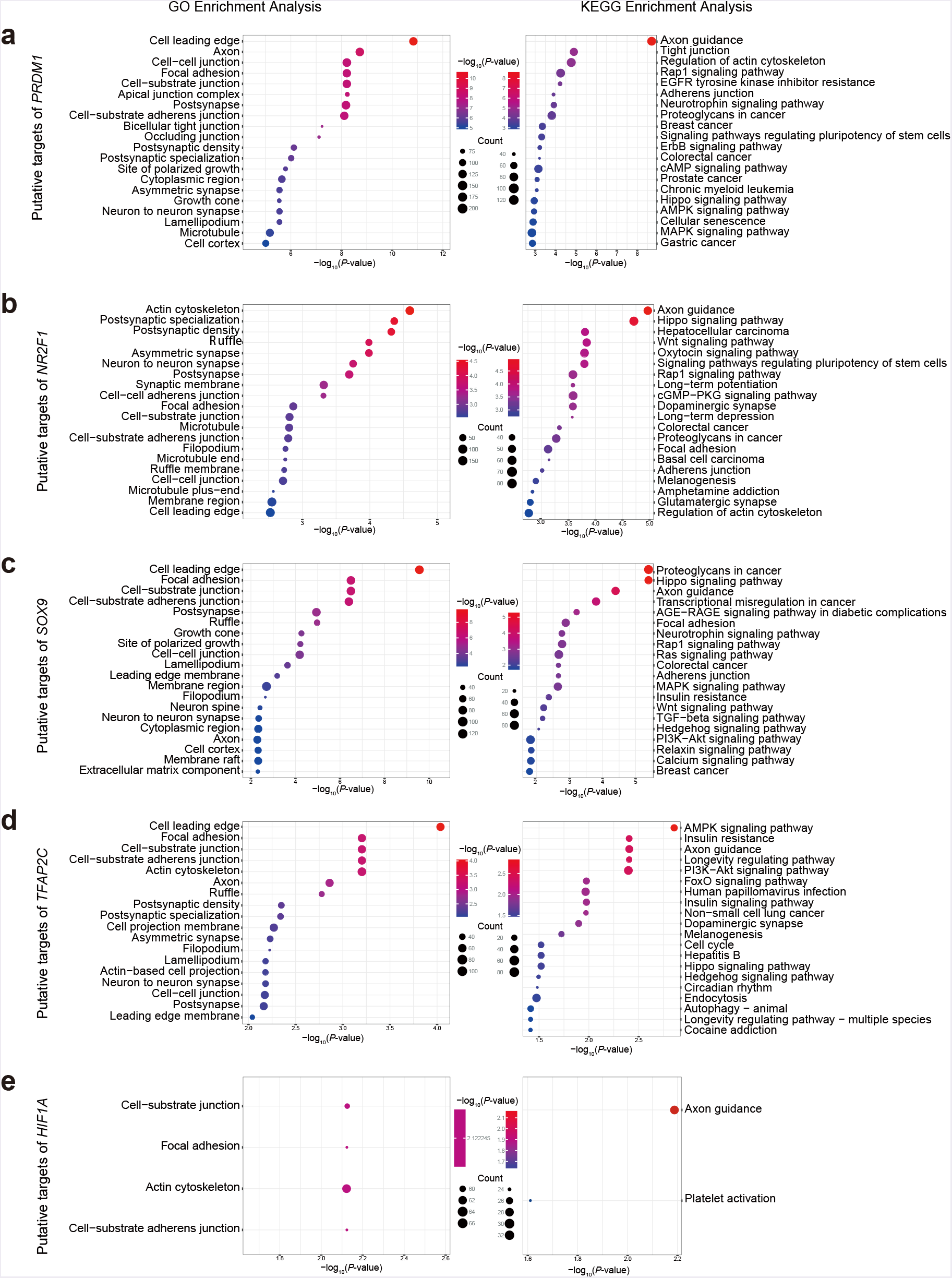
GO term and KEGG enrichment analysis of selected transcription factors (TFs) targets. GO term and KEGG enrichment analysis for putative targets of *PRDM1* (**a**), *NR2F1* (**b**), *SOX9* (**c**), *TFAP2C* (**d**) and *HIF1A* (**e**), adjusted P-value ≤ 0.05.

**Additional file 14: figure S14.**
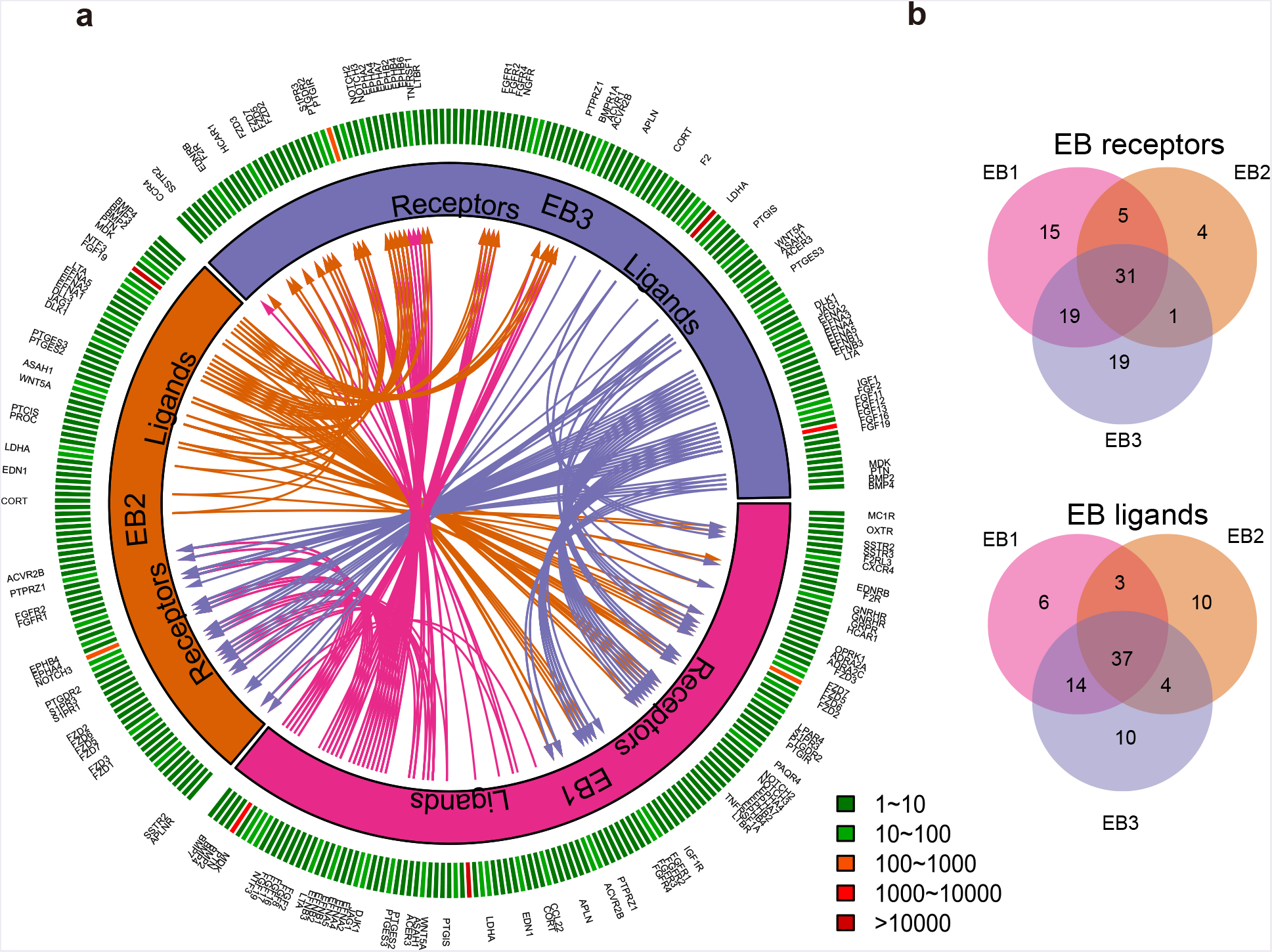
Putative signaling between expressed receptors and their ligands in EB subsets. **a** The inner layer compartments represent different cell subpopulations (EB1, EB2 and EB3). The outer layer indicates the expression profiles of ligands and receptors expressed in each cell subset, with low expressed molecular in green color while high expressed ones in red color. Arrows indicate putative interactions between ligands and receptors among cell subsets. **b** Venn plot showing the overlapping of ligands and receptors among cellular subpopulations.

**Additional file 15: figure S15.**
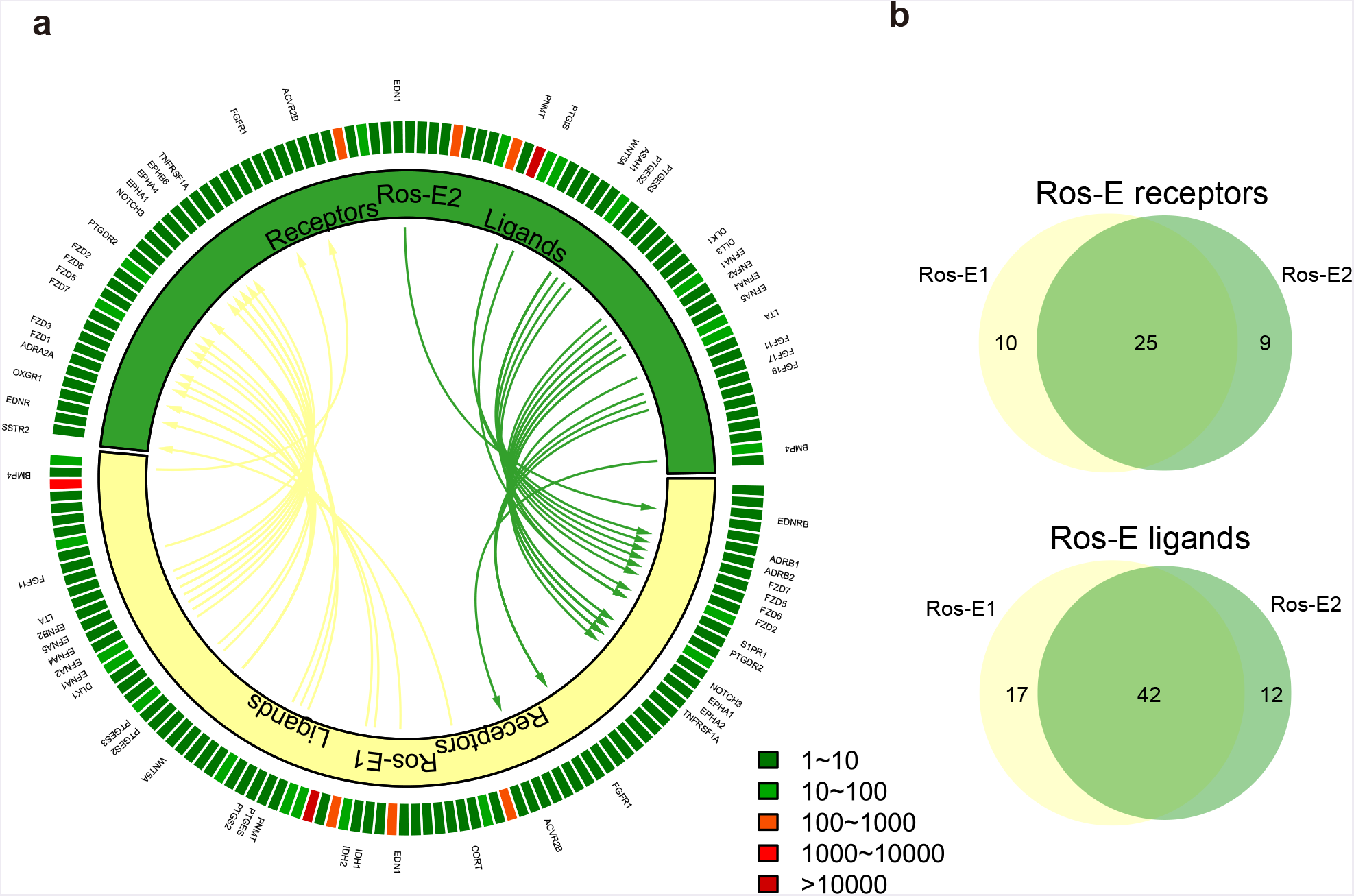
Putative signaling between expressed receptors and their ligands in Ros-E subsets. **a** The inner layer compartments represent different cell subpopulations (Ros-E1 and Ros-E2). The outer layer indicates the expression profiles of ligands and receptors expressed in each cell subset, with low expressed molecular in green color while high expressed ones in red color. Arrows indicate putative interactions between ligands and receptors among cell subsets. **b** Venn plot showing the overlapping of ligands and receptors among cellular subpopulations.

**Additional file 16: figure S16.**
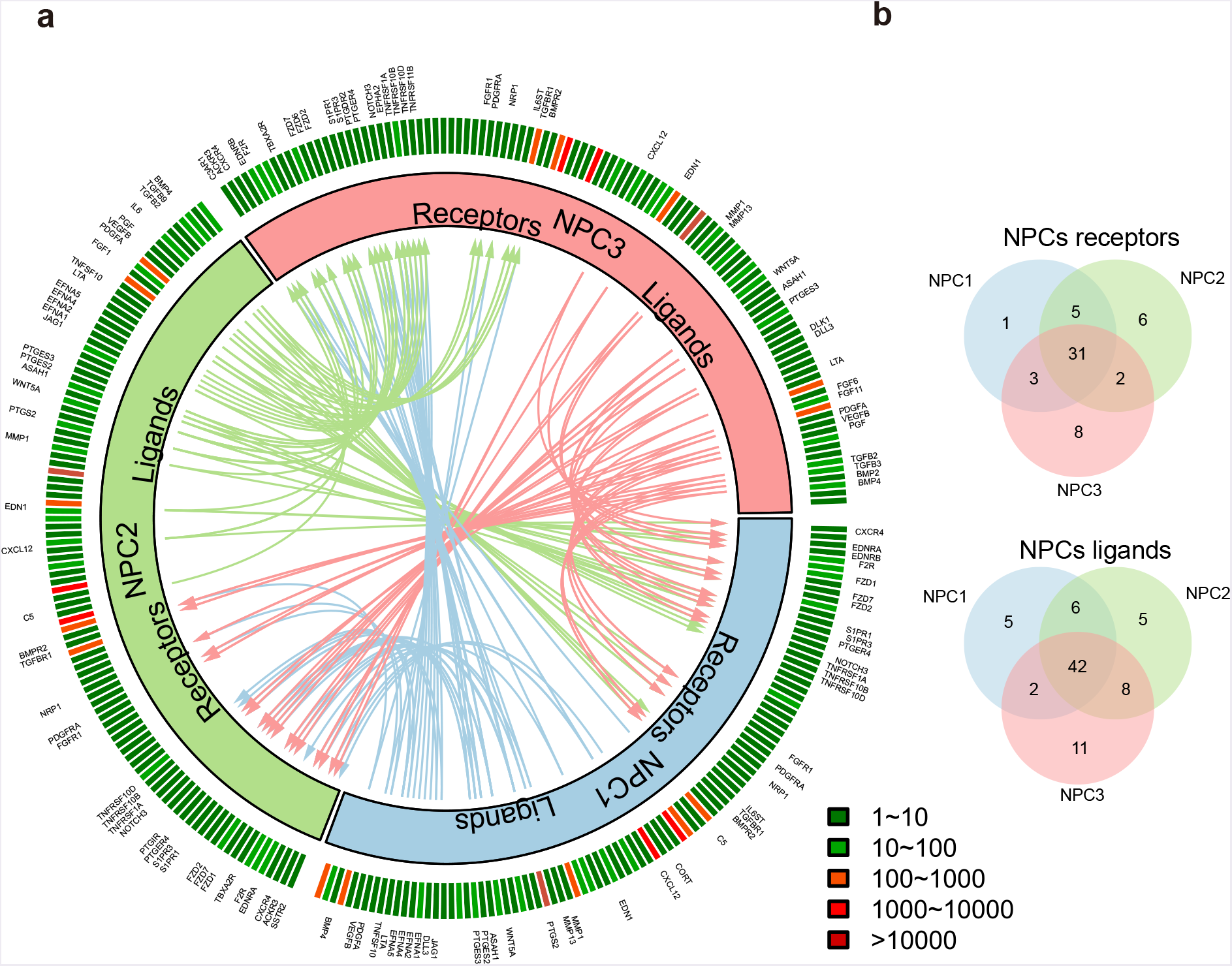
Putative signaling between expressed receptors and their ligands in NPC subsets. **a** The inner layer compartments represent different cell subpopulations (NPC1, NPC2 and NPC3). The outer layer indicates the expression profiles of ligands and receptors expressed in each cell subset, with low expressed molecular in green color while high expressed ones in red color. Arrows indicate putative interactions between ligands and receptors among cell subsets. **b** Venn plot showing the overlapping of ligands and receptors among cellular subpopulations.

**Additional file 17: figure S17.**
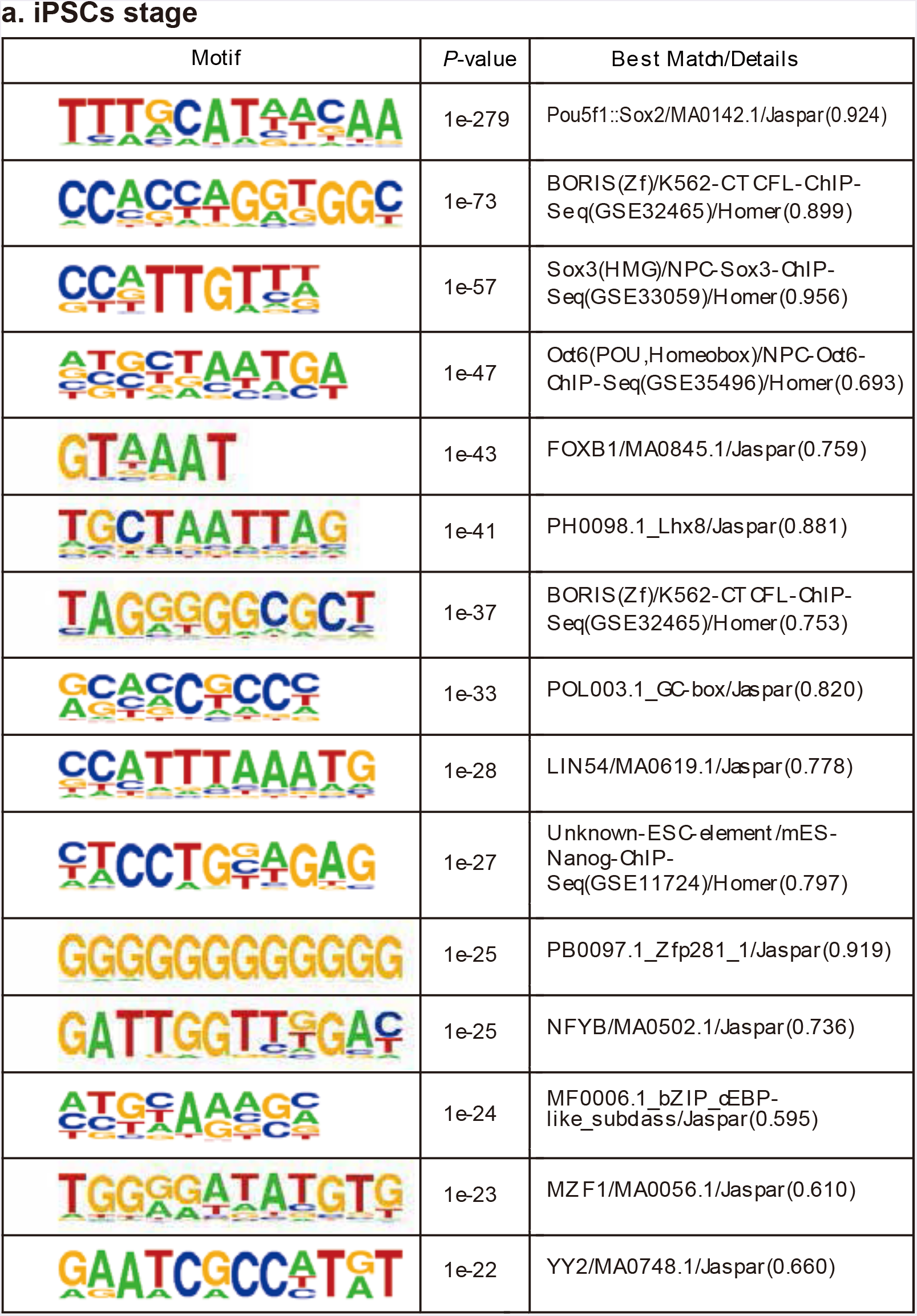
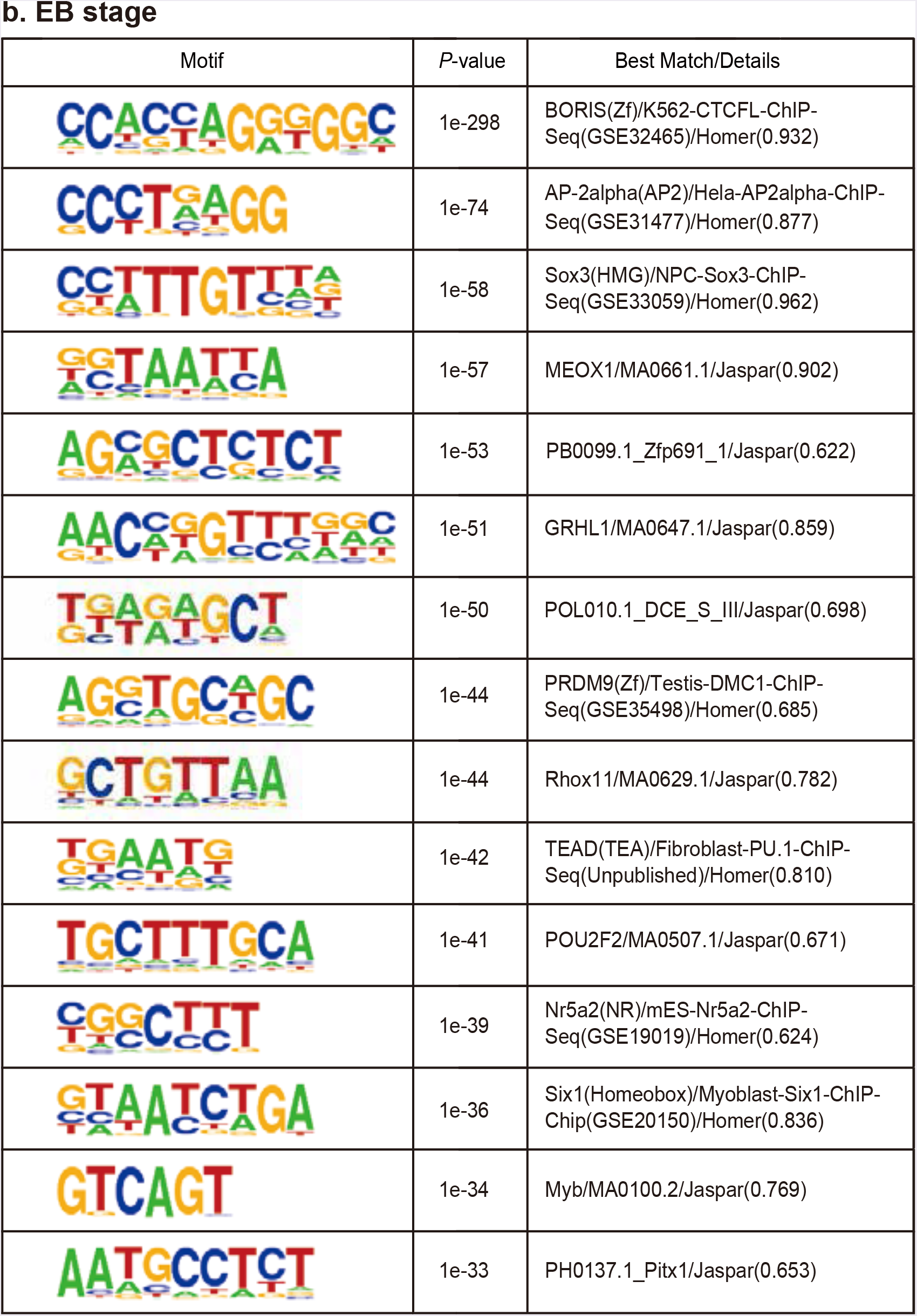
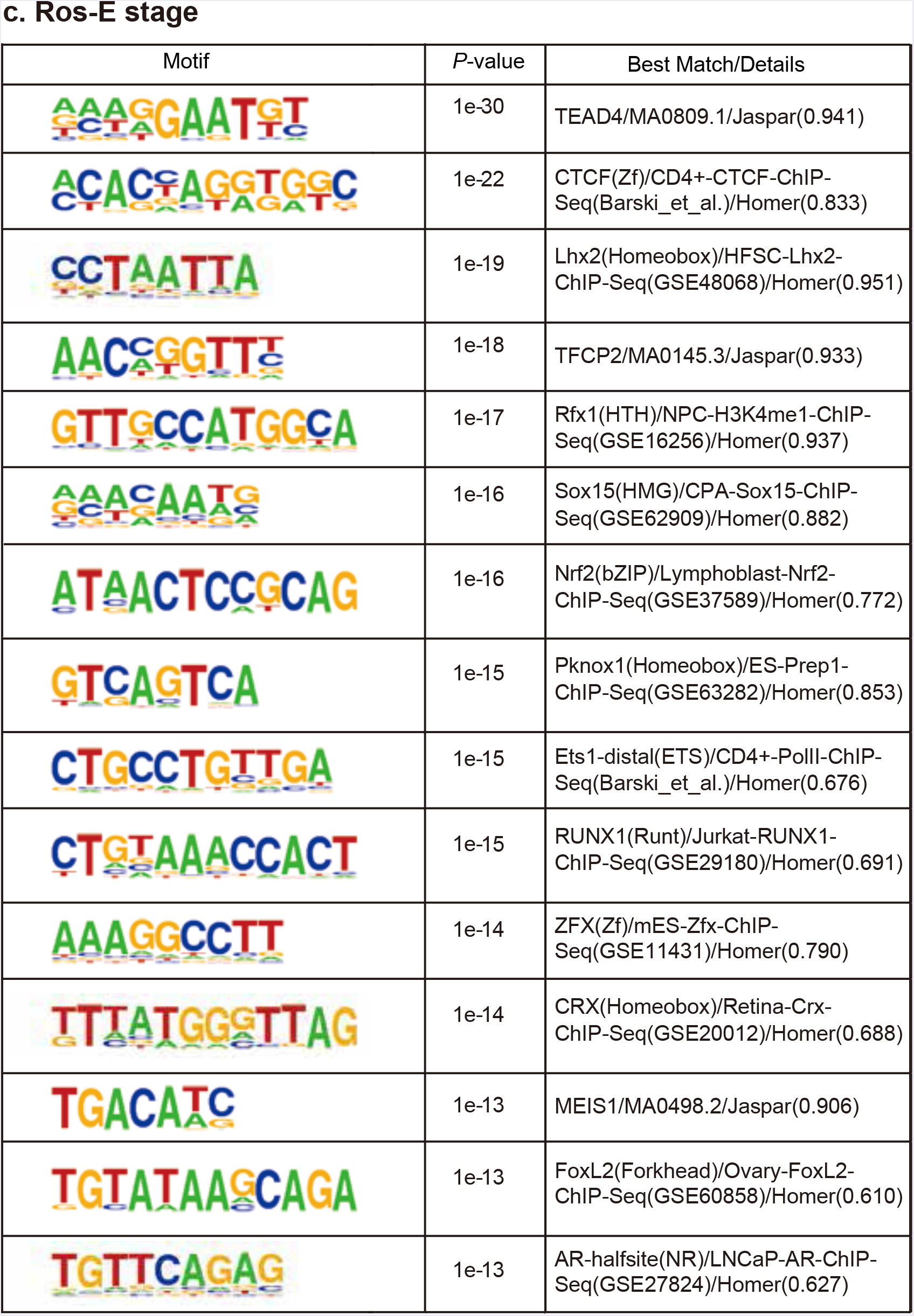
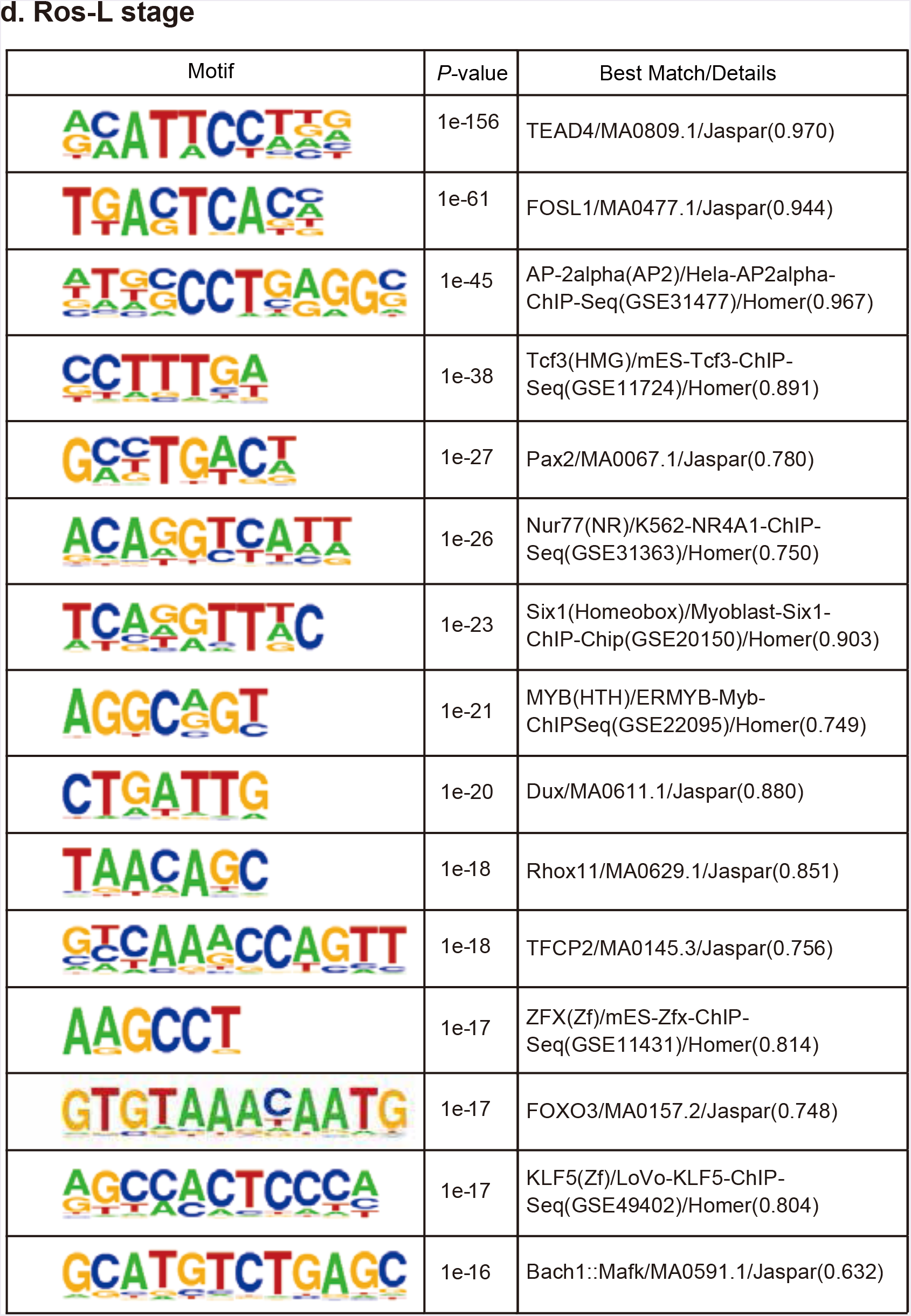
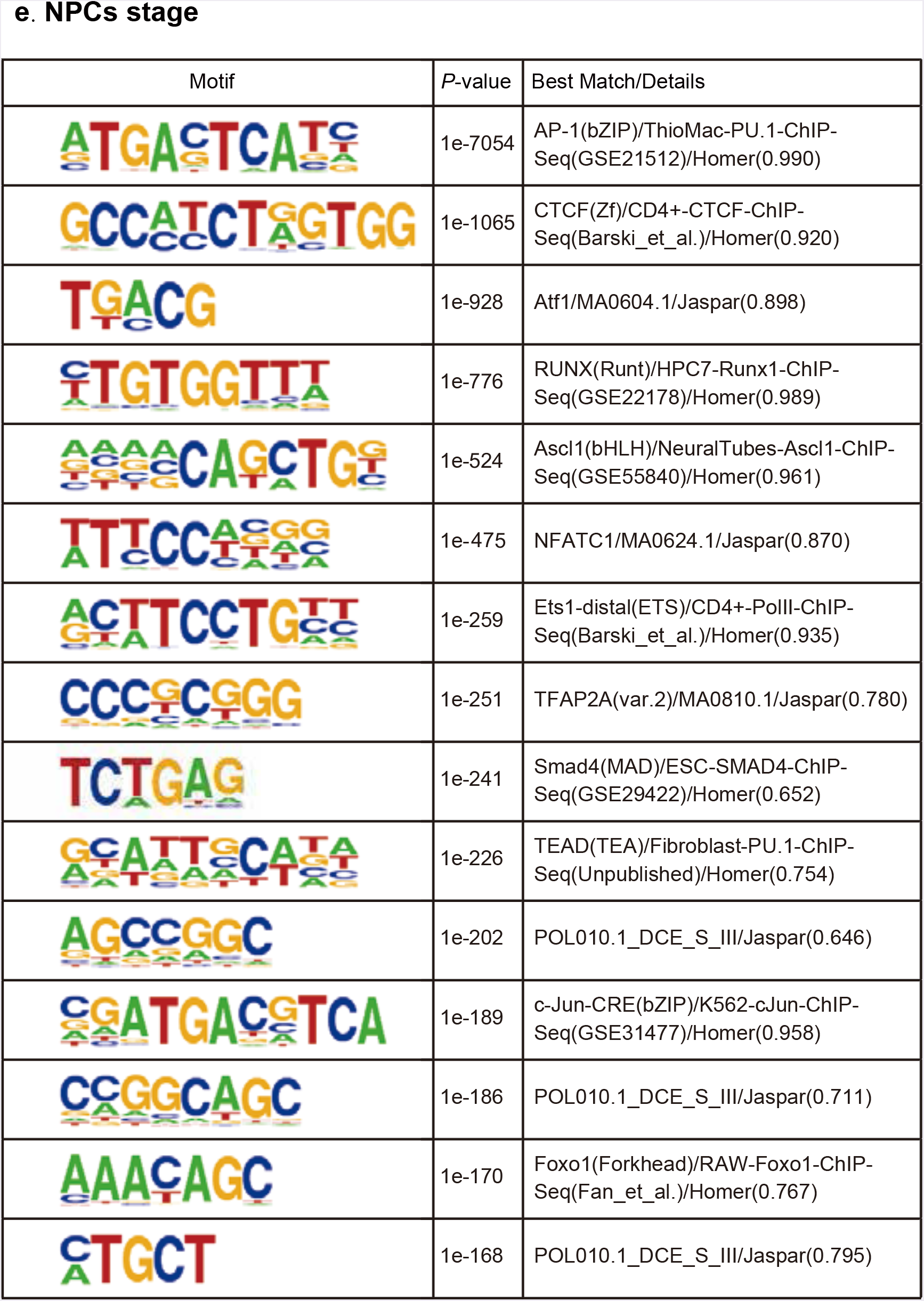
Transcription factor motifs enriched in stage specific peaks. Motifs enriched in stage specific ATAC peaks were listed in tables containing the following information: motif, P-value and best match/details for iPSCs (**a**), EB (**b**), Ros-E (**c**), Ros-L (**d**) and NPCs stage (**e**), respectively.

**Additional file 18: figure S18.**
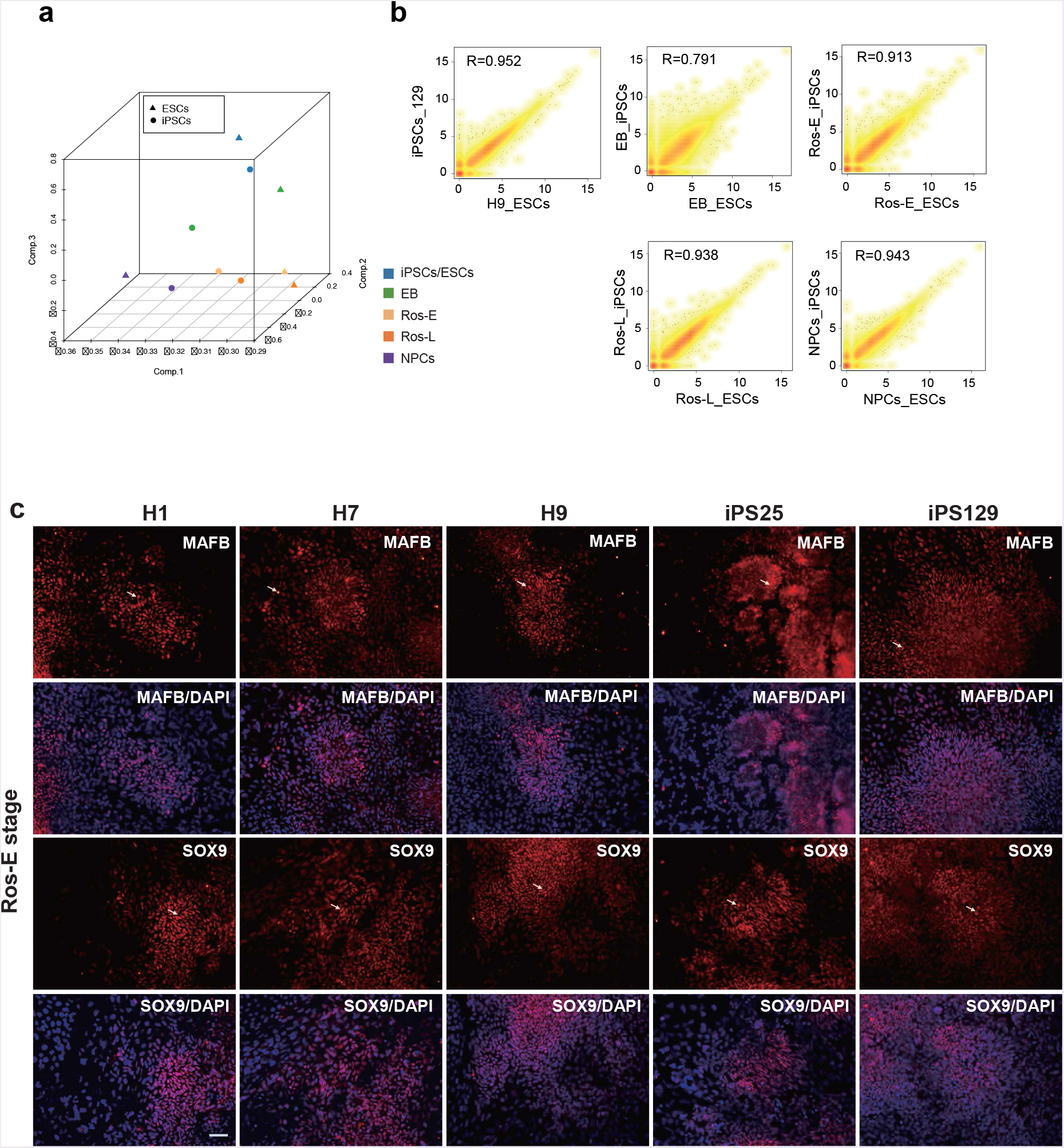
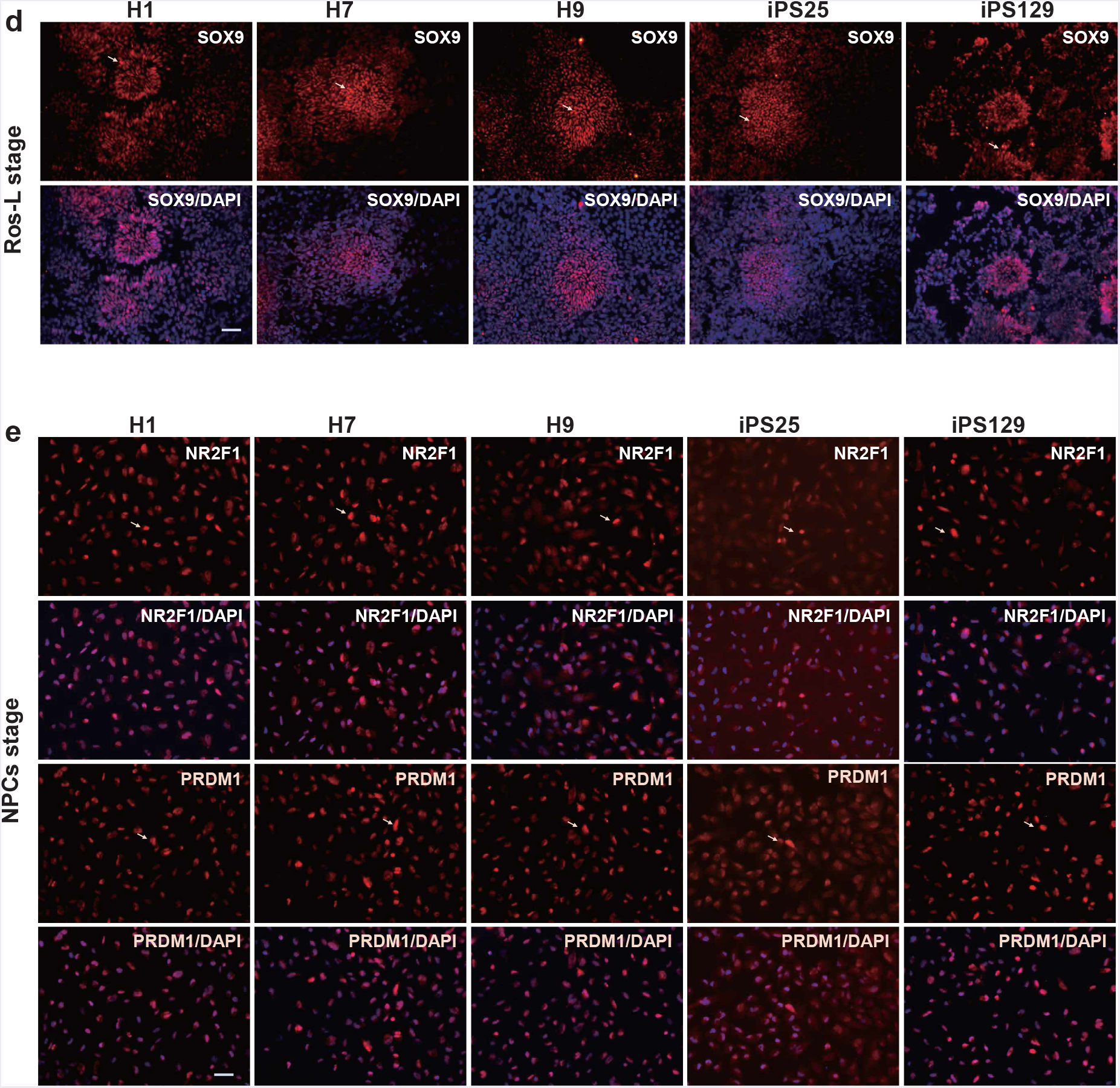
Validation of neural differentiation in different genetic background cell lines. **a** 3D PCA plot of the indicated cell stage derived from ESCs or iPSCs designated by colors and symbols. b The Pearson correlation coefficient between the corresponding cell stage derived from iPSCs and ESCs. **c, d, e** Immunostaining of MAFB and SOX9 at Ros-E stage (**c**), SOX9 at Ros-L stage (**d**), NR2F1 and PRDM1 at NPCs stage (**e**) across different genetic background cell lines (H1_ESCs, H7_ESCs, H9_ESCs, iPS25 and iPS129). Scale bar represents 50 pm.

**Additional file 19: Table S1** TFs differentially expressed among neighbouring cell subsets.

**Additional fie 20 Table S2** Putative targets of selected regulators.

**Additional file 21 Table S3** Subpopulations interaction networks.

**Additional file 22 Table S4** Differentially expressed receptors and ligands among Ros-L subpopulations.

## Availability of data and materials

The detailed protocol of neural differentiation and bioinformatics pipeline was available in protocol. io (DOI: dx.doi.org/10.17504/protocols.io.ntrdem6 and DOI: dx.doi.org/10.17504/protocols.io.ntpdemn). The sequencing raw data were deposited on NCBI SRA with the accession number SRP155759.

## Authors’ contributions

X.X. and Z.G. conceived and designed the project. Z.S., D.C., Q.W., S.W. and Q.D. conducted the majority of experiments and data analysis. L.W., X.D., S.W. and J.Z. performed computational analyses and prepared figures. C.L. participated in validation experiments and assisted with figure preparation for revision. D.Z., X.C. and F.C. contributed to sample collection. X.X., Z.G. and H.Y. supervised the project. X.L. contributed to the design of the revision and jointly supervised the validation work. Z.S., D.C., Q.W., Z.G. and X.X. prepared the manuscript. S.Z., L.L. and J.L.F. contributed to the discussion and revision of the manuscript. All authors read and approved the final manuscript.

## Acknowledgements

We thank Tao Tan for support with antibody, Shiping Liu for bioinformatics help, Guibo Li for technical help with preparation of single cell RNA-seq libraries and other members of Cell and Developmental Biology Lab for discussions and support. This work was supported by Shenzhen Engineering Laboratory for Innovative Molecular Diagnostics [grant number DRC-SZ [20,16] 884] funded by Development and Reform Commission of Shenzhen Municipality; and Shenzhen Key Laboratory of Neurogenomics (CXB201108250094A) funded by Science, Technology and Innovation Commission of Shenzhen Municipality. Dongsheng Chen is supported by China Postdoctoral Science Foundation (grant number 2017M622795).

